# Actin and AnkJ reciprocally regulate the catalytic state of the *Legionella* effector Ceg14

**DOI:** 10.64898/2026.09.15.751733

**Authors:** Dmitry Shatskiy, Marloes S. Eerelman, Siewert J. Marrink, Gunnar N. Schroeder, Alexander Belyy

## Abstract

Ceg14 is a *Legionella pneumophila* effector whose enzymatic activity is activated by host actin and inhibited by the cognate metaeffector AnkJ. Recent studies established that Ceg14 cleaves ATP to AMP and pyrophosphate and can transfer AMP to 3-phosphoglycerate, but how actin and AnkJ regulate these catalytic states remained unclear. Here, we show that Ceg14 is selectively activated by G-actin and determine the cryo-EM structure of the Ceg14-actin-AnkJ complex. Structure-guided mutagenesis identifies two actin-binding elements, a C-terminal anchor and an extended sensor region, that are required for Ceg14 activation and toxicity. Structural modelling and molecular dynamics simulations reveal that the catalytic center is assembled from three spatially distinct components: an N-terminal nucleoside-binding region, a phosphate-coordinating region within the actin sensor, and the catalytic triad. G-actin constrains the sensor and thereby organizes the phosphate-binding component of the active site. AnkJ binds at a distinct surface and allosterically repositions the N-terminal domain, disrupting nucleoside coordination while leaving part of the catalytic machinery intact. Consistent with this mechanism, AnkJ strongly suppresses AMP production and 3-phosphoglycerate-stimulated activity, yet the ternary complex retains weak Ceg14-dependent ATP-to-ADP activity. Thus, our results show that actin and AnkJ do not simply switch Ceg14 on and off, but reciprocally remodel its multipart catalytic center to determine its catalytic state and reaction output.

## Introduction

*Legionella pneumophila* is an opportunistic intracellular pathogen and the major causative agent of Legionnaires’ disease, a severe form of pneumonia (McDade et al., 1977; Fraser et al., 1977). In natural environments, *L. pneumophila* replicates within diverse protozoan hosts, whereas human infection occurs following inhalation of contaminated aerosols and subsequent uptake of the bacteria by alveolar macrophages (Boamah et al., 2017; Liu and Shin, 2019). Intracellular replication critically depends on the Dot/Icm type IV secretion system, which translocates more than 300 bacterial effector proteins into the host cytoplasm. Collectively, these effectors reprogram membrane trafficking, signaling, metabolism and other cellular processes to establish the *Legionella*-containing vacuole (LCV), a replication-permissive compartment that avoids canonical endolysosomal maturation (Isberg et al., 2009; Lockwood et al., 2022). The activities of some effectors are controlled by host factors but also by other translocated proteins, termed metaeffectors, providing an additional regulatory layer that can spatially or temporally constrain potentially detrimental effector activities (Joseph et al., 2021; Lockwood et al., 2022).

The actin cytoskeleton represents an important target of *Legionella* effectors because of its central roles in phagocytosis, membrane trafficking and organelle organization. *Legionella* produces five effectors that affect cytoskeleton interacting directly with actin. VipA and RavH bind monomeric actin and promotes filament nucleation, MavH associates with F-actin and promotes membrane-dependent actin assembly, RavK proteolytically cleaves actin, and RavJ catalyzes the covalent crosslinking of actin to members of the motin protein family (Franco et al., 2012; Zhang et al., 2026; Zhang et al., 2023; Liu et al., 2017; Liu et al., 2025). Moreover, actin can also serve as a host-derived regulator of effector activity as previously demonstrated for AMPylase LnaB and the related effector Ceg14 (Wang et al., 2024; He et al., 2025).

Ceg14 (Lpg0437; also known as SidL) was initially identified as an inhibitor of host protein synthesis (Fontana et al., 2011). Expression of Ceg14 is highly toxic in yeast, where it alters budding and actin organization, while recombinant Ceg14 inhibits actin polymerization in vitro (Guo et al., 2014).More recently, Ceg14 was shown to possess a conserved S-H-xxx-E motif, which upon binding of actin efficiently catalyzes the conversion of ATP to AMP and pyrophosphate, reducing cellular ATP levels (He et al., 2025). Ceg14 is regulated by the metaeffector AnkJ (Lpg0436; also known as LegA11), which directly engages with Ceg14 in a high-affinity 1:1 complex and strongly suppresses its ATP-consuming and translation-inhibiting activity (He et al., 2025; Machtens et al., 2026; Guan et al., 2026). These observations have established a model in which host actin functions as an activator of Ceg14, whereas AnkJ acts as its cognate inhibitor.

The biochemical consequences of Ceg14 activity, however, appear to extend beyond ATP depletion. Recent work identified the central glycolytic intermediate 3-phosphoglycerate (3PG) as a substrate of Ceg14 and showed that the enzyme transfers AMP from ATP to 3PG to generate the previously undescribed metabolite 2-AMP-3-phosphoglycerate (2-AMP-3PG) (Black et al., 2026). Production of 2-AMP-3PG in mammalian cells is accompanied by depletion of 3PG, disruption of glycolytic metabolism and inhibition of mTORC1 signaling, providing a mechanistic connection between Ceg14 activity and the previously observed inhibition of host translation. Importantly, 2-AMP-3PG is also produced during *L. pneumophila* infection of macrophages, supporting the physiological relevance of this reaction (Black et al., 2026). Ceg14 can therefore couple ATP cleavage to chemically distinct outcomes, either releasing AMP and pyrophosphate or transferring the AMP moiety to a metabolic substrate. How these catalytic states are established and how actin and AnkJ regulate the Ceg14 enzymatic activity has remained unclear.

Here, we determine the cryo-EM structure of the Ceg14-actin-AnkJ complex and use it to dissect the molecular basis of Ceg14 activation and metaeffector-mediated regulation. By combining structural analysis with targeted mutagenesis, biochemical activity measurements, yeast toxicity assays and molecular dynamics (MD) simulations, we define the interactions and conformational changes that distinguish different functional states of Ceg14 and control its catalytic output. We find that AnkJ does not simply abolish Ceg14 activity. Instead, AnkJ alters the catalytic state of the enzyme, suppressing AMP production while promoting ADP formation. Altogether, our findings provide a mechanistic framework for the regulation of Ceg14 by host actin and its cognate metaeffector AnkJ.

## Results

### Ceg14 is activated by G-actin and modulated by AnkJ

Actin exists in a dynamic equilibrium between monomeric globular actin (G-actin) and polymeric filamentous actin (F-actin). While purified actin can be maintained predominantly in its monomeric state under low-ionic-strength conditions, increasing the ionic strength to conditions resembling the intracellular environment strongly favors polymerization and shifts this equilibrium towards F-actin (Carlier, 1990). G- and F-actin can be experimentally stabilized using small molecules: phalloidin (PHD) strongly stabilizes F-actin (Estes et al., 1981; Pospich et al., 2020), whereas Latrunculin A (LatA) binds G-actin and prevents its incorporation into filaments (Yarmola et al., 2000). Distinguishing between these two states in biochemical assays is important because many actin-targeting toxins preferentially recognize either G- or F-actin. For example, the actin ADP-ribosylating toxins C2, iota toxin and CDT (Tsuge et al., 2008; Gülke et al., 2001; Aktories et al., 2011) modify monomeric G-actin but not F-actin, whereas TcART, ExoY, VopV, SipA and other bacterial factors preferentially interact with actin in its filamentous state (Belyy et al., 2022; Belyy et al., 2021; Hiyoshi et al., 2011; Niedzialkowska et al., 2024).

Ceg14 was previously shown to hydrolyze ATP to AMP and pyrophosphate (PPi) in an actin-dependent manner (He et al., 2025). However, because enzymatic activity was measured under ionic conditions that permit spontaneous filament assembly, the actin state responsible for this activation was not established. We therefore compared recombinantly produced Ceg14 in the presence of phalloidin-stabilized F-actin and LatA-sequestered G-actin. Previous studies monitored Ceg14 nucleotide conversion primarily by qualitative or semi-quantitative HPLC or thin-layer chromatography approaches (He et al., 2025; Black et al., 2026). To quantitatively determine Ceg14 activity, we adapted a colorimetric assay based on detection of free orthophosphate and analyzed reactions in parallel with and without inorganic pyrophosphatase. This allowed us to distinguish ATP-to-ADP hydrolysis, which directly releases phosphate, from ATP-to-AMP conversion, which produces PPi and becomes detectable after its conversion into two molecules of orthophosphate. In the presence of LatA-sequestered G-actin, Ceg14 showed robust AMP production of 173 ± 30 nmol AMP min^-1^ mg^-1^, whereas we detected minimal activity with phalloidin-stabilized F-actin and no measurable activity in the absence of actin (Fig. 1A). The previously reported catalytically inactive mutant Ceg14 E575A (He et al., 2025) showed no detectable activity in the presence of G-actin. These results establish G-actin as the specific activator of Ceg14 and provide a quantitative measure of its actin-dependent ATP-to-AMP activity.

**Figure 1.**
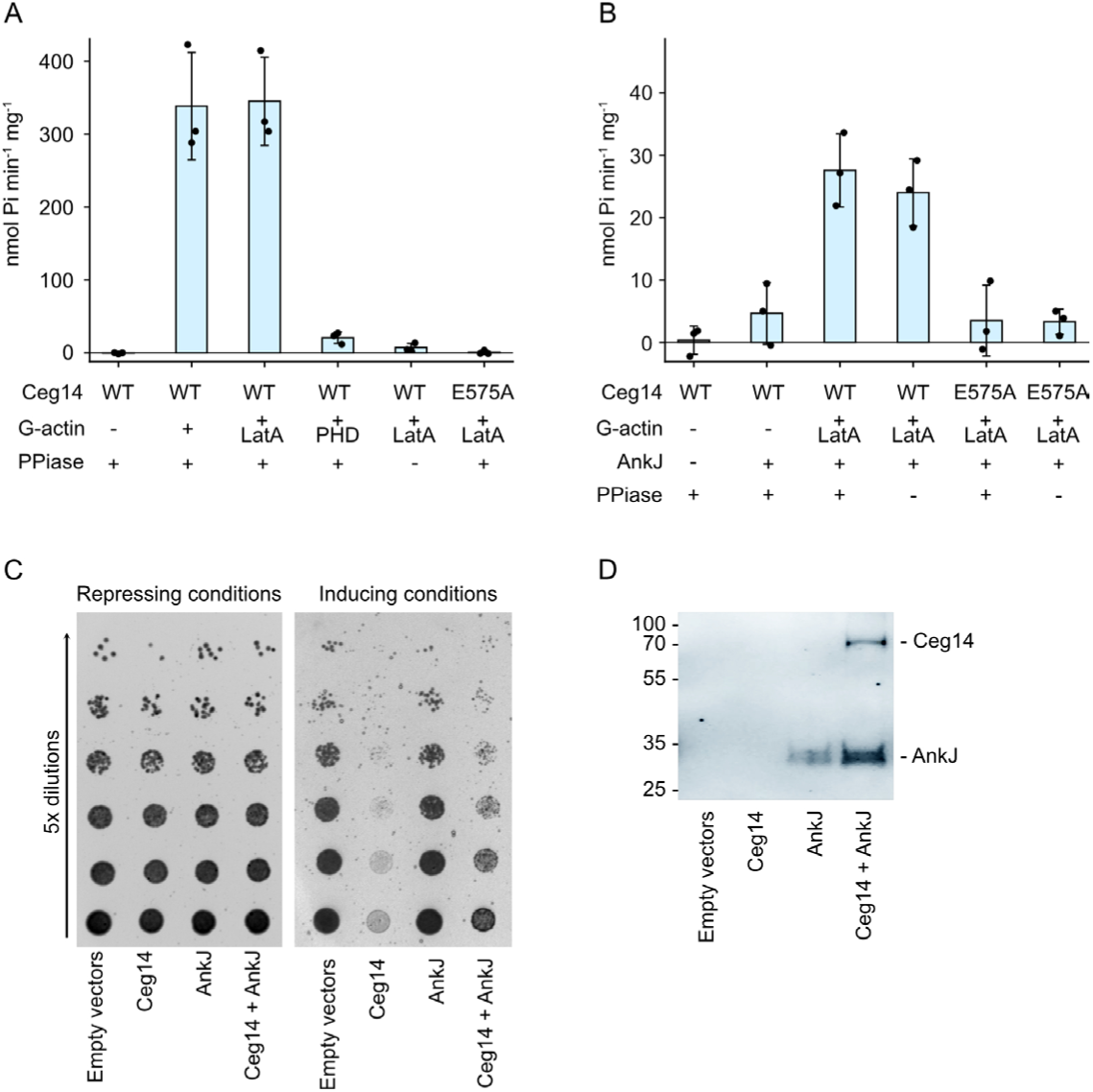
Ceg14 is activated by G-actin and modulated by AnkJ. (A, B) ATP hydrolysis activity of 0.4 µg Ceg14 in the presence of 3 µg actin (stabilized by equimolar concentration of latrunculin, LatA, or phalloidin, PHD) and 2.2 µg AnkJ measured using a malachite green phosphate assay. In the presence of inorganic pyrophosphatase (PPiase), the assay detects ATP conversion to both ADP and AMP, whereas in the absence of PPiase only ATP-to-ADP hydrolysis is detected because pyrophosphate does not react with malachite green. ATP hydrolysis by actin is strongly coupled to filament assembly; therefore, sequestration of G-actin by LatA prevents polymerization-associated ATP hydrolysis. Under our assay conditions, the residual ATPase activity of monomeric G-actin was negligible and did not measurably contribute to the phosphate signal. The data are presented as mean values, the error bars correspond to standard deviations of three independent experiments. (C) Yeast cells expressing Ceg14 from a galactose-inducible promoter and AnkJ from the strong TEF1 constitutive promoter were serially diluted, spotted on solid medium with 2% glucose (repressing conditions) or with 2% galactose and 0.05% glucose (inducing conditions), and incubated for 2 days at 30 °C. (D) Western blot analysis showing the production of Ceg14 and AnkJ, both fused to an N-terminal Myc tag in yeast cultured under inducing conditions. C, D representative of two independent experiments.

Ceg14 is regulated by its cognate metaeffector AnkJ, which directly binds the effector and has been reported to inhibit its activity (He et al., 2025; Machtens et al., 2026). We therefore quantified Ceg14 activity in the presence of both AnkJ and G-actin. In the absence of actin, the Ceg14-AnkJ complex showed almost no ATP hydrolysis (Fig. 1B). When we added G-actin, however, we observed a distinct change in catalytic output. Whereas Ceg14-actin produced predominantly AMP, the Ceg14-actin-AnkJ complex showed nearly no AMP production but ADP-producing activity of 24 ± 5 nmol ADP min^-1^ mg^-1^, approximately 7-fold lower than the AMP-producing activity of Ceg14-actin. The catalytically inactive Ceg14 E575A variant in complex with actin and AnkJ showed background level of activity, confirming that the detected ADP production depends on the Ceg14 catalytic center. Thus, AnkJ does not simply abolish Ceg14 catalysis. Instead, it strongly reduces overall activity while shifting the catalytic output from AMP towards ADP production.

*Saccharomyces cerevisiae* has been widely used as a heterologous model to study *Legionella* effectors, as many of their cellular targets and the pathways they manipulate are conserved in yeast (Campodonico et al., 2005; Heidtman et al., 2009; Urbanus et al., 2016). Yeast is particularly suitable for studying the actin-dependent activity of Ceg14 because *S. cerevisiae* contains a single conventional actin encoded by ACT1, avoiding the complexity introduced by multiple cytoplasmic actin isoforms in mammalian cells (Belmont and Drubin, 2001). We examined whether AnkJ suppresses Ceg14 toxicity in yeast. We expressed Ceg14 from a galactose-inducible promoter, while expressing AnkJ from a strong constitutive promoter to ensure that AnkJ was already present when Ceg14 production was induced. Expression of Ceg14 caused a severe growth defect, whereas co-expression with AnkJ reduced Ceg14 toxicity (Fig. 1C). However, AnkJ did not completely restore yeast growth. Western blot analysis showed that AnkJ accumulated to higher levels than Ceg14 under these conditions (Fig. 1D), ruling out insufficient AnkJ expression as cause of the incomplete rescue. Similar incomplete suppression of Ceg14 toxicity was reported previously when both genes were expressed from galactose-inducible promoters (Urbanus et al., 2016; He et al., 2025).

The persistence of Ceg14-dependent toxicity is notable given the very high affinity of the interaction between Ceg14 and AnkJ, which form a 1:1 complex with a reported dissociation constant of approximately 1.8 nM (Machtens et al., 2026). Moreover, other *Legionella* metaeffectors can completely neutralize the activity of their cognate effectors; for example, SidD reverses SidM-mediated AMPylation of Rab1 and completely rescues SidM-induced toxicity in yeast (Tan and Luo, 2011), and MesI forms a high-affinity complex with SidI and fully suppresses SidI-mediated toxicity (Joseph et al., 2020). Together with our biochemical measurements, these observations support a model in which AnkJ strongly reduces, but does not completely abolish, the enzymatic activity of Ceg14.

### Cryo-EM structure of the Ceg14-actin-AnkJ complex

To understand how actin activates Ceg14 and how AnkJ regulates its activity, we aimed to determine cryo-EM structures of both the Ceg14-actin and Ceg14-actin-AnkJ complexes. Despite extensive screening of different constructs and sample conditions, we were unable to find intact Ceg14-actin particles suitable for structural analysis. In contrast, the Ceg14-actin-AnkJ complex yielded well-defined particles, allowing us to determine its structure at an overall resolution of 3.6 Å (Fig. 2A, Supplementary Table S1, Supplementary Figure S1, S2).

**Figure 2.**
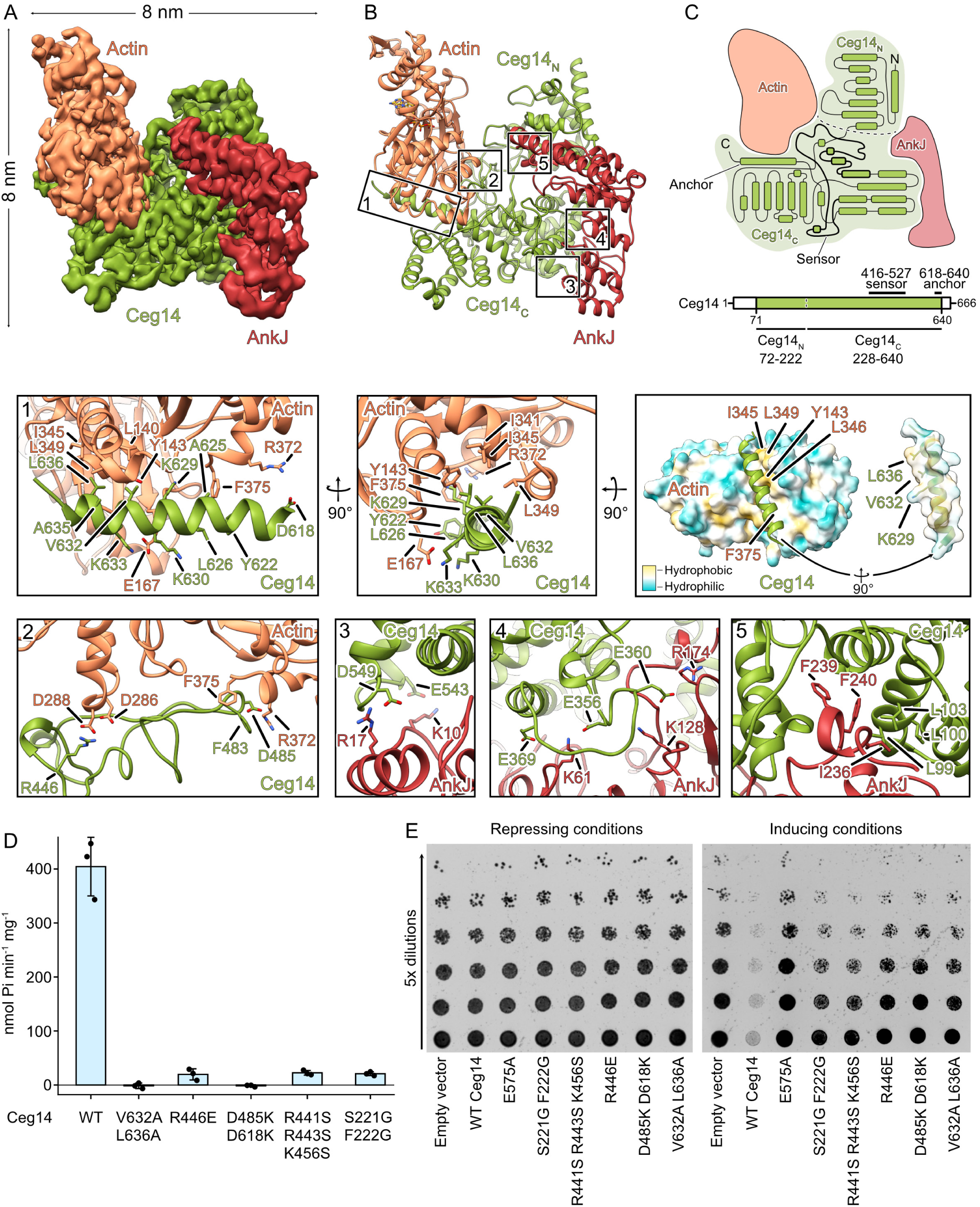
Cryo-EM structure of the Ceg14-actin-AnkJ complex. 3.6 Å Cryo-EM reconstruction (A) and the structure (B) of the Ceg14-actin-AnkJ complex. Ceg14, actin and AnkJ are in green, orange and red, respectively. (C) Schematic illustration of the domain organization and secondary structure of Ceg14. (D) ATP hydrolysis activity of Ceg14 and its mutants in the presence of LatA-stabilized G-actin, measured after incubation with PPiase. The data are presented as mean values, the error bars correspond to standard deviations of three independent experiments. (E) Yeast cells expressing Ceg14 or its mutants from the galactose-inducible promoter were serially diluted and spotted on the medium with 2% glucose (repressing conditions) or with 2% galactose and 0.05% glucose (inducing conditions), and incubated for 2 days at 30 °C. D, E representative of two independent experiments.

The Ceg14-actin-AnkJ complex adopts a compact 1:1:1 assembly with overall dimensions of approximately 8 x 8 x 6 nm (Fig. 2A-C). Ceg14 forms the central scaffold of the complex, with actin and AnkJ bound to opposite sides of the effector. Ceg14 itself is an elongated, predominantly alpha-helical protein composed of two major globular domains connected by a short linker. We did not observe density corresponding to the extreme N-terminus of Ceg14, suggesting that this part of the protein is flexible. The resolved density begins at residue 72, which forms a compact globular domain, termed the N-terminal domain, or Ceg14_N_, extending to residue 222. This domain consists of eight alpha-helices connected by short loops. A short linker connects Ceg14_N_ to the larger C-terminal domain, Ceg14_C_, comprising residues 228-640. Ceg14_C_ is likewise almost entirely alpha-helical and contains no beta-sheet elements. It terminates in a long C-terminal helix, with the final 26 residues not resolved, again suggesting conformational flexibility.

Actin adopts a conformation closely resembling previously determined structures of native G-actin, with a C-alpha RMSD of 1.03 Å relative to native G-actin (Supplementary Figure S3A) (PDB 3HBT; Wang et al., 2010). As expected for monomeric actin, subdomain II is poorly resolved, and we observed no density for the DNase I-binding loop (D-loop), consistent with the pronounced flexibility of this region in G-actin. AnkJ is composed predominantly of alpha-helical elements, with 15 alpha-helices and a single two-stranded beta-sheet connected by intervening loops (Fig. 2B). AnkJ binds to the side of Ceg14 opposite the major actin-binding surface and does not make direct contacts with G-actin. Thus, actin and AnkJ can simultaneously engage spatially distinct surfaces of Ceg14 within the ternary complex.

AnkJ forms an extensive interaction interface with Ceg14 involving both its N-terminal and C-terminal domains. The C-terminal domain of Ceg14 engages the inner surface of the AnkJ ankyrin-repeat region by establishing numerous electrostatic interactions, including contacts involving Ceg14 E356, E360, E369, E543 and D549 (Fig. 2B). The N-terminal domain of Ceg14 provides an additional hydrophobic interaction surface. In particular, a hydrophobic patch formed by Ceg14 L99, L100 and L103 packs against AnkJ residues I236, F239 and F240. We next asked whether AnkJ engages Ceg14 in the same manner in the presence and absence of actin. Interestingly, during processing of the Ceg14-actin-AnkJ dataset, we identified a distinct population of smaller particles corresponding to the Ceg14-AnkJ complex (Supplementary Figure S1 and S2). We processed these particles independently and obtained a reconstruction at an overall resolution of 3.6 Å. The resulting Ceg14-AnkJ structure closely matched the Ceg14-AnkJ part of our ternary complex, C-alpha RMSD of 0.88 Å, and was highly similar to the previously reported crystal structure, C-alpha RMSD of 1.3 Å (Machtens et al., 2026) (Supplementary Figure S3 B and C). This demonstrates that actin has no impact on the contact network between AnkJ and Ceg14.

Ceg14 forms two extensive interaction interfaces with G-actin, both mediated by its C-terminal domain (Fig. 2B). The first interface is formed by the C-terminal helix of Ceg14, which contacts actin subdomains I and III through an extensive network of hydrophobic and electrostatic interactions. Ceg14 residues Y622, L626, V632, A635 and L636 form a large hydrophobic surface that packs against a complementary hydrophobic region of actin formed by L140, Y143, I341, I345, L349 and F375 (Fig. 2B, subpanel 1). Remarkably, K629 of Ceg14 is positioned within this otherwise highly hydrophobic interface. Inspection of the structure shows that its side-chain amino group directly contacts the C-terminal carboxylate of actin residue F375. The interface is further stabilized by salt bridges between Ceg14 K633 and actin E167, and between Ceg14 D618 and actin R372. Given the size and extensive nature of this interaction surface, we refer to the C-terminal helix of Ceg14 as the actin anchor. To test the functional importance of this region, we purified a Ceg14 V632A/L636A mutant designed to disrupt the hydrophobic interaction with actin. In our *in vitro* activity assay, these substitutions completely abolished actin-dependent ATP hydrolysis, indicating a loss of Ceg14 activation (Fig. 2D). Consistently, the V632A/L636A variant showed strongly reduced toxicity in yeast compared with wild-type Ceg14 (Fig. 2E). Together, these results identify the C-terminal anchor as a major actin-binding element of Ceg14 that is essential for its activation and toxicity.

The second major interaction interface, which we propose to call sensor, is formed by Ceg14 residues 416-527. Structurally, this region contains relatively little regular secondary structure: only 23 of its 112 residues form alpha helices, comprising three short helices and one longer 13-residue helix, while the remainder adopts an extended loop-rich conformation. Unlike the compact interaction surface formed by the C-terminal anchor, this sensor engages actin over a broad complementary surface through contacts distributed throughout its sequence. Within this extended interface, several individual contacts contribute to the interaction with G-actin. Two features are particularly prominent. Ceg14 R446 protrudes and forms electrostatic contacts with actin D286 and D288 (Fig. 2B, subpanel 2). Another residue of Ceg14, D485, is positioned close to the anchor and, together with D618, interacts with actin R372. To test the functional importance of the sensor, we produced and purified the Ceg14 R446E and D485K/D618K mutants. Both variants showed strongly reduced ATP-hydrolyzing activity in the presence of G-actin (Fig. 2D). Similarly, both mutants were substantially less toxic than wild-type Ceg14 in yeast (Fig. 2E). Together, these results identify the sensor as a second essential actin-binding element of Ceg14 and show that interactions mediated by this region are required for efficient actin-dependent activation and cytotoxicity.

### Mechanism of activation and inhibition of Ceg14

Having established that G-actin activates Ceg14 whereas AnkJ suppresses its AMP-producing activity, and having defined their interaction interfaces, we next sought to understand how these regulators control the catalytic center of Ceg14. Because we could not obtain an experimental structure of the Ceg14-actin complex, we generated a structural model of Ceg14 bound to actin and ATP using Boltz-2 (Passaro et al., 2025), with our experimentally determined Ceg14-actin-AnkJ structure supplied as a template. In this model, ATP occupies a catalytic cleft formed at the interface between the N-terminal and C-terminal domains of Ceg14 (Fig. 3A). The catalytic center can be divided into three functional components. First, the nucleotide base is accommodated by F222, C408, N412, H529 and F570, while S221 is positioned to interact with the 3’-hydroxyl group of ribose. Second, the phosphate groups are coordinated by the positively charged residues R441, R443 and K456. Third, H571 is positioned close to the alpha-phosphate and forms the catalytic triad together with S527 and E575. Importantly, these components originate from distinct structural regions of Ceg14 (Fig. 3E). S221 and F222 are located in Ceg14_N_, the phosphate-coordinating residues R441, R443 and K456 lie within the actin sensor, whereas the remaining nucleotide-binding residues and the catalytic triad reside in Ceg14_C_. Formation of a catalytically competent active site therefore requires the precise spatial organization of several structurally distinct regions of Ceg14, suggesting a mechanism by which ligand binding at sites distant from the catalytic cleft could regulate enzyme activity.

**Figure 3.**
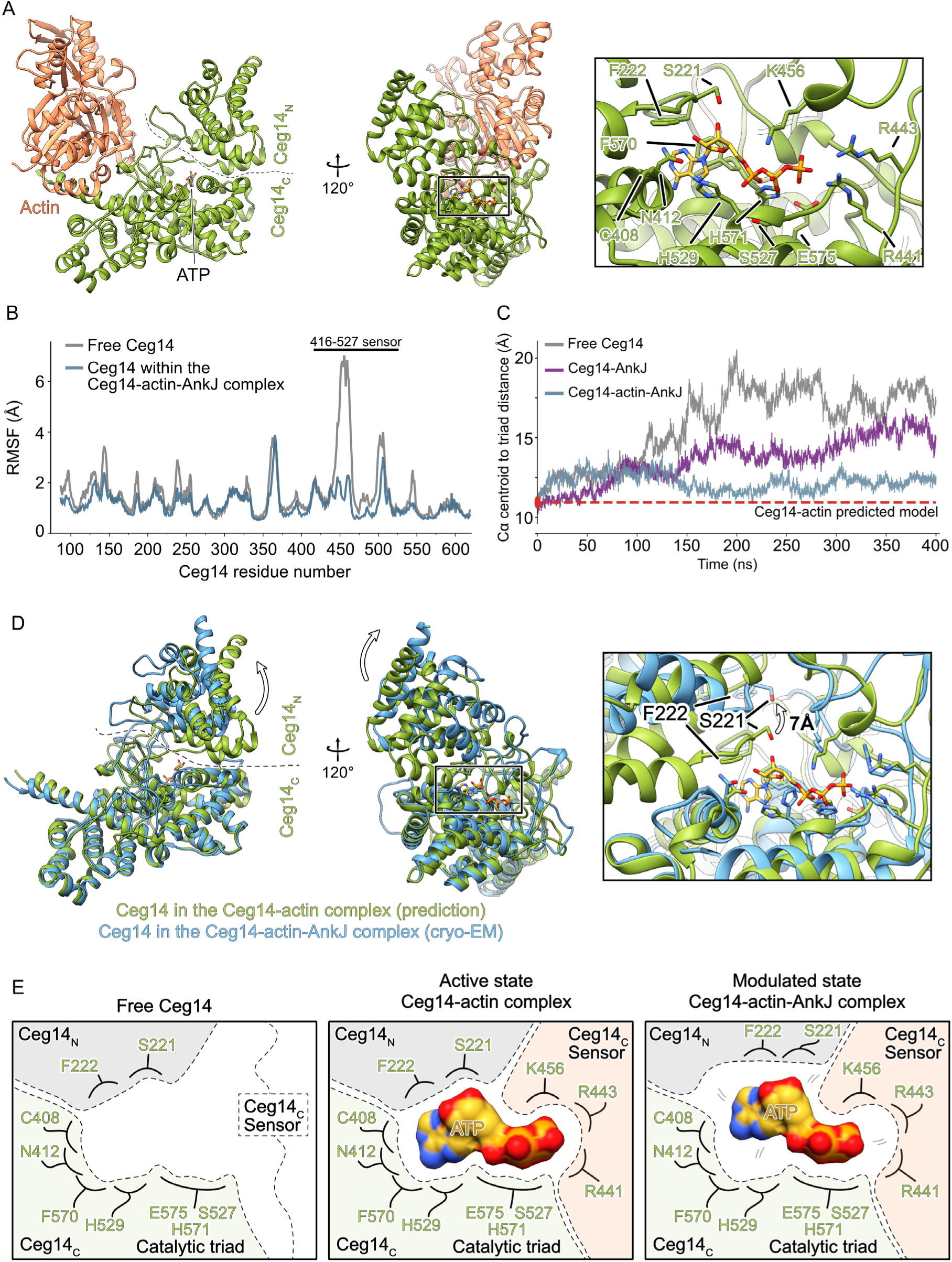
Mechanism of activation and inhibition of Ceg14. (A) Boltz-2 prediction of the Ceg14-actin-ATP complex. (B) Per-residue RMSF of Ceg14 in the free form and within the Ceg14-actin-AnkJ complex within the last 100 ns of the all-atom MD simulation of 400 ns. The curve represents the average values from 3 independent simulations. (C) Distance between C-alpha of R441, R443 and K456 to the catalytic triad during all-atom MD simulations. The curve represents the average values from 3 independent simulations. Red dotted line represents the distance between these residues in the Ceg14-actin model. (D) Alignment between Ceg14 models from the Ceg14-actin prediction (green) and the Ceg14-actin-AnkJ structure (blue). Actin and AnkJ are removed for clarity. (E) Schematic organization of the Ceg14 catalytic center in the free Ceg14, as well as in the Ceg14-actin and Ceg14-actin-AnkJ complexes.

We next asked how G-actin promotes this organization. To examine the conformational dynamics of Ceg14 in the absence of its binding partners, we removed actin and AnkJ from our experimentally determined structure and performed all-atom molecular dynamics simulations of Ceg14 alone. Most regions of Ceg14 showed comparatively limited conformational variability both in the free protein and in the Ceg14-actin-AnkJ complex (Fig. 3B). In contrast, residues 430-465 displayed pronounced flexibility in free Ceg14, reaching RMSF values of up to 7 Å compared with approximately 2 Å in the ternary complex. Consistent with this observation, the same region is not resolved in our Ceg14-AnkJ structure (Supplementary Figure 3B), further indicating that it remains flexible in the absence of actin. Strikingly, this mobile segment forms part of the actin sensor and contains both the actin-contacting residue R446 and the phosphate-coordinating residues R441, R443 and K456. R446 directly engages actin D286 and D288, placing the actin-binding site immediately adjacent to the residues responsible for coordinating the ATP phosphate groups. This arrangement provides a direct structural link between actin recognition and assembly of the catalytic center. R441, R443 and K456 remained within 12 Å to the catalytic triad in the simulations in the presence of actin, similarly to the distance in the Ceg14-actin complex, while drifting further away in free Ceg14, as well as in the Ceg14-AnkJ complex (Fig. 3C). We therefore propose that binding of G-actin constrains the flexible sensor and positions R441, R443 and K456 for productive ATP coordination, explaining why the Ceg14 R441S/R443S/K456S mutant lost detectable ATP-hydrolyzing activity in the presence of G-actin and showed strongly reduced toxicity in yeast (Fig. 2 D and E). Together, our structural, biochemical and simulation data support an allosteric activation mechanism in which G-actin organizes the sensor region and thereby completes the catalytic center of Ceg14.

We next asked how AnkJ inhibits Ceg14. Because AnkJ binds outside the catalytic cleft, its effect should be transmitted allosterically. To examine this mechanism, we compared Ceg14 in the predicted Ceg14-actin complex with the effector in our experimentally determined Ceg14-actin-AnkJ structure. The C-terminal domain adopts a highly similar conformation in both states, whereas AnkJ binding is associated with a substantial displacement of the N-terminal domain, with local shifts of up to 10 Å (Fig. 3D). This rearrangement affects the three components of the catalytic center differently. The phosphate-coordinating residues and the catalytic triad remain in nearly identical positions, whereas the N-terminal domain contributing S221 and F222 to the nucleoside-binding region, moves away from the catalytic cleft. Consequently, ATP can no longer be accommodated in the geometry observed in the predicted active state. Consistent with an essential role of this region in catalysis, the Ceg14 S221G/F222G mutant showed little ATP-hydrolyzing activity in the presence of G-actin and strongly reduced toxicity in yeast (Fig. 2D and E). The functional importance of these residues is further supported by their complete conservation among the Ceg14 sequences analyzed (Supplementary Fig. S4A).

Importantly, the AnkJ-induced displacement of the N-terminal domain selectively disrupts the nucleoside-binding region without dismantling the entire catalytic center. The phosphate groups can still be accommodated by the positively charged residues of the sensor, while the catalytic triad remains structurally intact. However, displacement of S221 and F222 prevents the adenine and ribose moieties from adopting the orientation observed in the predicted active state and thereby disrupts productive positioning of ATP relative to the catalytic histidine. At the same time, loss of tight coordination by the nucleoside-binding region may allow ATP to remain bound in a less constrained and more conformationally heterogeneous configuration. In such a state, the catalytic histidine may gain access to alternative phosphate positions and promote reactions that are disfavored in the fully assembled active site. This provides a possible explanation for the ATP-to-ADP activity that we observed for the Ceg14-actin-AnkJ complex *in vitro*. We therefore propose that AnkJ suppresses AMP-producing activity by selectively disrupting nucleotide-binding geometry while leaving a partially assembled catalytic center capable of alternative chemistry.

The three-component organization of the catalytic center therefore provides a mechanistic explanation for how actin and AnkJ can regulate Ceg14 while remaining simultaneously bound to the effector (Fig. 3E). Actin stabilizes the sensor region and thereby organizes the phosphate-coordinating component of the active site, whereas AnkJ acts at a spatially distinct surface repositioning the N-terminal domain and disrupting the nucleoside-binding component. Thus, the two regulators do not compete for Ceg14 binding but instead exert opposing allosteric effects on different elements of the same catalytic center. Their simultaneous binding maintains Ceg14 in a distinct regulatory state in which part of the catalytic machinery remains assembled, AMP-producing activity is suppressed, and a low level of alternative ADP-producing activity becomes possible.

### AnkJ suppresses the 3PG-dependent activity of Ceg14

During preparation of this manuscript, Ceg14 was reported to function as an adenylyltransferase that uses 3-phosphoglycerate (3PG) as an AMP acceptor, generating 2-AMP-3PG (Black et al., 2026). We therefore asked whether the presence of 3PG influences Ceg14 catalytic activity in our *in vitro* system. Using the malachite green-based assay described above, we measured ATP hydrolysis in the presence or absence of 3PG. Addition of 3PG at a concentration equimolar to ATP increased the rate of ATP hydrolysis approximately twofold, consistent with 3PG acting as a more efficient acceptor substrate for Ceg14 (Fig. 4). We next asked whether 3PG could similarly stimulate ATP hydrolysis by the Ceg14-actin-AnkJ complex, which retains weak ADP-producing activity but no detectable AMP-producing activity. In contrast to Ceg14-actin, addition of 3PG did not increase the activity of the ternary complex, arguing against efficient transfer of an ADP-containing group to 3PG under these conditions.

**Figure 4.**
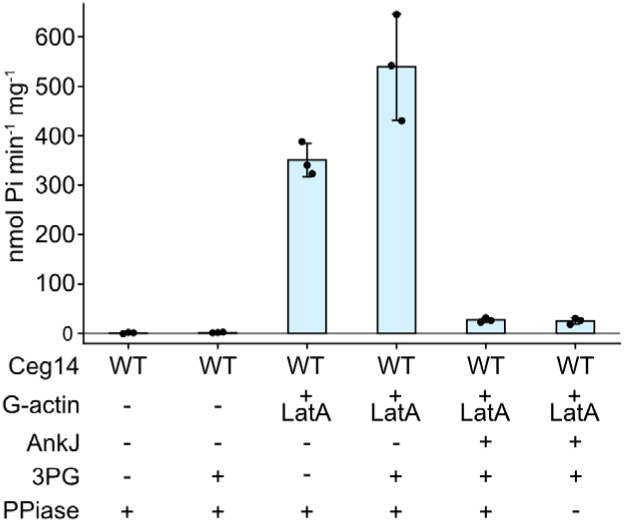
AnkJ suppresses the 3PG-dependent activity of Ceg14. ATP hydrolysis activity of 0.4 µg Ceg14 in the presence of 3 µg LatA-stabilized actin, 2.2 µg AnkJ and 1 mM 3PG using a malachite green phosphate assay. The data are presented as mean values, the error bars correspond to standard deviations of three independent experiments.

This result is consistent with our structural model of AnkJ-mediated regulation. Productive transfer of AMP to 3PG is expected to require precise positioning of ATP, the acceptor substrate and the catalytic residues within the active site. In the actin-activated state, all three components of the Ceg14 catalytic center are assembled, providing the constrained geometry required for this reaction. AnkJ binding displaces the nucleoside-coordinating region and disrupts this geometry while leaving part of the catalytic machinery intact. This altered state may still permit the weak, less constrained ATP-to-ADP reaction observed *in vitro* but appears incompatible with the precise substrate organization required for efficient transfer to 3PG. Thus, AnkJ not only suppresses AMP production by Ceg14 but also prevents stimulation of its catalytic activity by the physiological acceptor substrate 3PG.

## Discussion

Our study provides a refined mechanistic framework for the reciprocal regulation of the catalytic state of Ceg14 by host actin and the metaeffector AnkJ. We identify G-actin as the activating form of actin and show that Ceg14 recognizes it through two interaction regions: a C-terminal anchor and an extended sensor. The use of actin as an activating host factor may be particularly advantageous for *Legionella*, which replicates in phylogenetically diverse protozoan hosts and can opportunistically infect mammalian macrophages. Actin is one of the most highly conserved eukaryotic proteins (Pollard, 2016), and our sequence comparison shows striking conservation of the Ceg14-binding surface across the analyzed species (Supplementary Fig. S4B). Ceg14 activation may therefore be insensitive to host-species variation, in contrast to more host-adapted effectors such as the *Legionella* amylase LamA, whose activity has distinct consequences in its natural amoebal hosts and in mammalian cells (Price et al., 2020).

Our structure helps reconcile observations made before the enzymatic activity of Ceg14 was known. Previous work showed that Ceg14 toxicity in yeast was suppressed by profilin overexpression and identified the non-toxic Ceg14 variants G234V and T623N, which had lost the ability to inhibit actin polymerization (Guo et al., 2014). Our structure places T623 directly within the C-terminal actin anchor, whereas G234 lies close to the junction between the N- and C-terminal domains, where its substitution may interfere with the conformational coupling required for activation. Interestingly, the Ceg14-binding surface on actin overlaps with that recognized by profilin (Funk et al., 2019). However, rather than excluding Ceg14 activity, very large excesses of profilin are required to substantially suppress Ceg14-mediated ATP hydrolysis *in vitro*: Ceg14 remains active even at profilin:actin ratios far above stoichiometric levels (He et al., 2025). Because a major fraction of cellular G-actin is normally associated with profilin (Funk et al., 2019), this observation suggests that profilin-bound actin may constitute a physiologically relevant reservoir of G-actin for activating Ceg14. Ceg14 may therefore have evolved not to rely on a rare pool of completely free actin, but instead to efficiently capture G-actin from its normal cellular complexes.

The organization of the Ceg14 catalytic center also explains several biochemical observations from previous studies. Earlier mutational analyses established the importance of the residues S527, H571 and E575 and identified N412 and R443 as essential for ATP hydrolysis, with substitutions of G528, H529 and Y565 also reducing activity (He et al., 2025). Our structural model places these residues within a multipart catalytic center assembled from the N-terminal domain, the actin-responsive sensor and the C-terminal domain. Interestingly, we propose that S221 is positioned to interact with the 3’-hydroxyl group of the ribose. Because this interaction does not require the 2’-hydroxyl group, the geometry is compatible with deoxyribose and provides a simple explanation for the previously reported actin-dependent conversion of dATP to dAMP (He et al., 2025). The nucleotide pocket also appears relatively permissive: although the base is surrounded by F222, C408, N412, H529 and F570, the predicted model does not reveal an obvious network of interactions uniquely specifying adenine. Thus, Ceg14 can accommodate nucleotide substrates other than ATP, reflecting its promiscuous nucleotide specificity.

A parallel structural study (Guan et al., 2026) independently reached a broadly compatible model of allosteric Ceg14 regulation. Their Ceg14-actin structure showed that actin binding reorganizes a flexible region they termed the “Lid”, largely overlapping our sensor, and induces movement of the N-terminal domain. They proposed that the N-terminal domain undergoes transitions between closed, intermediate and open conformations, with the open state positioning H571 for productive ATP cleavage. Conversely, AnkJ restricts these dynamics and traps Ceg14 in an intermediate conformation. Our results refine this model by linking these conformational changes to distinct components of the catalytic center. Actin engages the sensor immediately adjacent to R441, R443 and K456 and thereby organizes the phosphate-binding component, whereas AnkJ acts from the opposite surface to displace the N-terminal contribution to nucleoside coordination, including S221 and F222. Importantly, our biochemical data indicate that the AnkJ-bound state is not completely catalytically inactive: although AMP production is strongly suppressed, the Ceg14-actin-AnkJ complex retains weak Ceg14-dependent ATP-to-ADP activity. This differs from a strict inactive-state model and suggests instead that AnkJ changes the reaction geometry of Ceg14. Retention of the phosphate-coordinating region and catalytic triad while nucleoside coordination is disrupted could permit less constrained ATP positioning and thereby enable low-efficiency chemistry that is disfavored in the fully assembled active state. The incomplete suppression of Ceg14 toxicity by AnkJ in yeast is also consistent with AnkJ strongly attenuating rather than necessarily eliminating every catalytic state of the effector, although the contribution of residual ADP-producing activity to cellular toxicity remains to be established.

Ceg14 belongs to the S-HxxxE family of bacterial effectors, another well-characterized member of which is the *Legionella* effector LnaB. Like Ceg14, LnaB requires host actin for activity and uses ATP for AMP-transfer chemistry; however, LnaB functions as a phosphoryl-AMPylase that transfers AMP to phosphorylated substrates, including phosphoribosyl-ubiquitin (PR-Ub) and phosphorylated host proteins (Fu et al., 2024; Wang et al., 2024; Chen et al., 2026). Despite their related catalytic architecture and common dependence on actin, the mechanisms by which the two enzymes achieve a catalytically competent state appear distinct. In LnaB, binding of the acceptor substrate, such as PR-Ub, induces rearrangement of residues M217 and F256 and opens access to the ATP-binding channel, thereby stimulating catalysis. In Ceg14, productive ATP binding does not appear to require an analogous substrate-induced opening step. Instead, our data indicate that G-actin itself organizes the catalytic center by constraining the sensor region and positioning its phosphate-coordinating residues. Thus, related S-HxxxE enzymes use different conformational mechanisms to couple host-factor recognition to ATP-dependent chemistry. This distinction is particularly relevant in light of the recent identification of 3PG as a physiological AMP acceptor for Ceg14 (Black et al., 2026). Whereas 3PG further stimulated ATP turnover by the actin-activated enzyme, it did not stimulate the Ceg14-actin-AnkJ complex in our assays. These observations suggest that, unlike the acceptor-driven activation observed for LnaB, Ceg14 is first activated by G-actin to generate an AMP-transfer-competent catalytic center, while AnkJ subsequently disrupts this state and prevents modification of 3PG.

Together, our results in line with parallel studies (Guan et al., 2026; Black et al., 2026) support a model (Fig. 5) in which Ceg14 activity is determined by the assembly state of a multipart catalytic center. In free Ceg14, the actin sensor is conformationally flexible and the residues required for phosphate coordination are not stably organized. Binding of G-actin through the C-terminal anchor and sensor constrains this region and positions the phosphate-coordinating residues relative to the nucleoside-binding elements and the catalytic triad, generating a catalytically competent state that cleaves ATP at the alpha-phosphate and supports AMP transfer to 3PG. This conversion of 3PG into 2-AMP-3PG depletes a central glycolytic intermediate and thereby impairs host glycolytic metabolism. AnkJ can bind simultaneously on the opposite side of Ceg14 and does not displace actin. Instead, it allosterically repositions the N-terminal domain of Ceg14, moving S221 and F222 away from the catalytic cleft while leaving the sensor and catalytic triad assembled. This selectively disrupts the geometry required for AMP-producing chemistry and 3PG modification, while permitting a low level of alternative ATP-to-ADP activity. Thus, rather than functioning as a simple on/off switch, actin and AnkJ reciprocally remodel different components of the same catalytic center to determine the reaction output of Ceg14. Understanding when and where the Ceg14-actin-AnkJ complex forms during infection, and what physiological function this distinct catalytic state serves, will therefore be an important direction for future studies.

**Figure 5.**
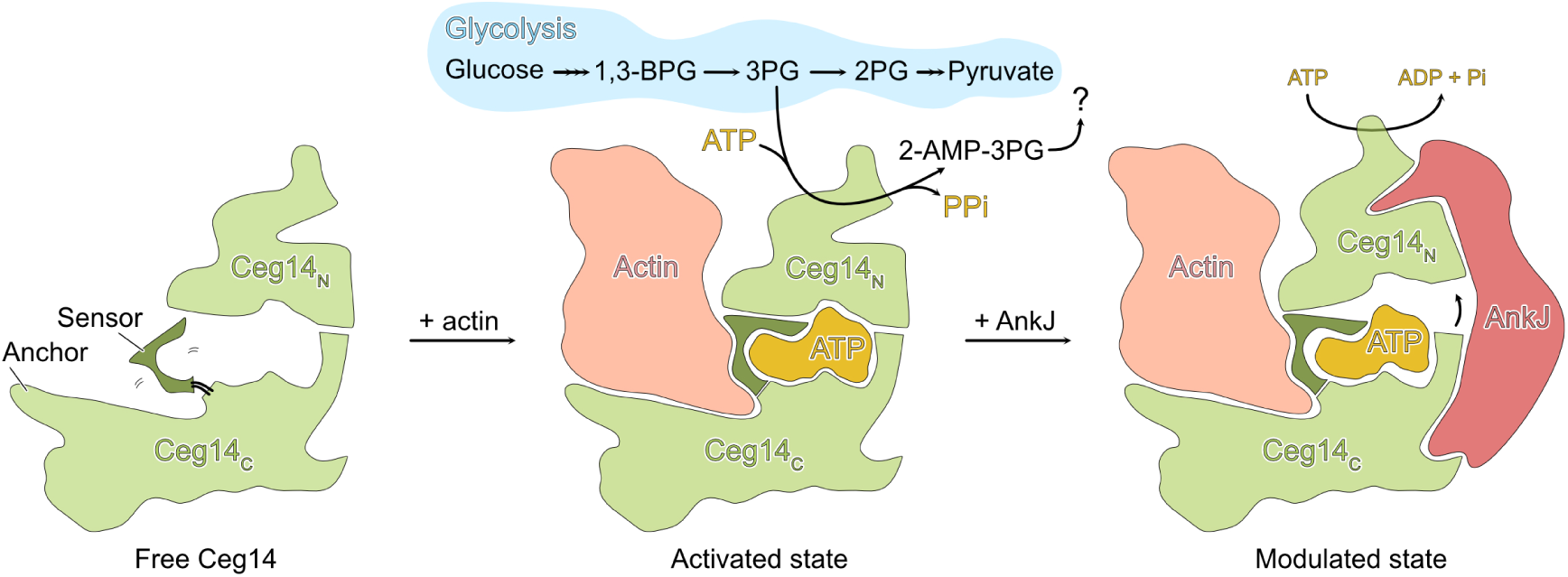
Schematic illustration of activation and inhibition of Ceg14.

## Materials and Methods

### Protein production and purification

The codon-optimized nucleotide sequence encoding Ceg14 (Uniprot Q5ZYD5) was synthesized by Twist Bioscience as gene fragments and cloned as a 6xHis-maltose-binding protein (MBP) fusion into pB137 (Belyy et al., 2018), a pET28a-derived expression vector, using the SacI and HindIII restriction sites. Initial attempts to produce full-length Ceg14 without MBP yielded insufficient soluble protein, whereas fusion to MBP substantially improved protein production and solubility and was therefore used throughout this study. AnkJ (Uniprot Q5ZYD6) was similarly synthesized and cloned into pET28a using the SacI and HindIII sites. Ceg14 mutants were generated by synthesizing DNA fragments carrying the desired substitutions and replacing the corresponding parts of the wild-type construct.

MBP-Ceg14 and AnkJ were produced in *Escherichia coli* BL21-CodonPlus(DE3)-RIPL cells following overnight induction at 22 °C with 0.02 mM IPTG. Cells were lysed in 20 mM Tris-HCl, pH 8.0, 150 mM NaCl, and the soluble fraction was applied to a Ni-IDA affinity column. The column was washed with lysis buffer, and bound proteins were eluted with the same buffer supplemented with 250 mM imidazole. Ceg14-containing fractions were immediately subjected to size-exclusion chromatography on a Superdex 200 Increase 10/300 GL column equilibrated in 20 mM Tris-HCl, pH 8.0, 150 mM NaCl. We found that freezing and thawing substantially reduced Ceg14 enzymatic activity; therefore, purified Ceg14 and its variants were stored on ice, used within one week of purification, and were never frozen. AnkJ-containing fractions were dialyzed against lysis buffer for 2-3 h, aliquoted, and stored at -20 °C. In contrast to Ceg14, AnkJ remained stable under these conditions, and its inhibitory activity did not measurably change during storage.

Rabbit skeletal muscle alpha-actin was purified as described previously (Belyy et al., 2021). Briefly, muscle acetone powder (Pel-Freez Biologicals) was resuspended in G-buffer containing 5 mM Tris-HCl, pH 7.5, 1 mM DTT, 0.2 mM CaCl_2_, and 0.5 mM ATP. The suspension was centrifuged for 30 min at 100,000 x g to remove insoluble material. The G-actin-containing supernatant was supplemented with MgCl2 and KCl to final concentrations of 2 mM and 100 mM, respectively, to induce actin polymerization. After 1 h at room temperature, KCl was added to a final concentration of 800 mM to dissociate actin-binding proteins, and F-actin was pelleted by centrifugation for 2 h at 100,000 x g. The F-actin pellet was subsequently dialyzed against G-buffer for 5 days to depolymerize the filaments. Purified G-actin was flash-frozen in liquid nitrogen and stored in small aliquots at -70 °C.

For enzymatic assays and cryo-EM sample preparation, freshly thawed alpha-actin was centrifuged at 120,000 x g for 20 min at 4 °C using a TLA-120.1 rotor, and the supernatant containing monomeric G-actin was collected. Where indicated, G-actin was mixed with an equimolar concentration of Latrunculin A. F-actin was generated by incubating G-actin for 2 h on ice in F-buffer containing 120 mM KCl, 20 mM Tris-HCl, pH 8.0, 2 mM MgCl2, 1 mM DTT, and 1 mM ATP. The resulting filaments were stabilized by addition of an equimolar concentration of phalloidin (Fisher Scientific).

### Yeast toxicity experiments

Ceg14 WT and its variants were amplified by PCR from the corresponding bacterial expression plasmids using the primers TATACTCGAGCAGAATCTGGATGAAATTCTTAAGAAATTAC and TATATAGCTAGCTCAGCAGCCATACACTGTTCC. The PCR products were digested with XhoI and NheI and cloned into pK87 (Belyy et al., 2015), a derivative of YEpGal555 carrying a galactose-inducible promoter. AnkJ was amplified from the corresponding pET28-based expression plasmid using the primers TATACTCGAGATCAAGATGGGCCGATCGGAAATGAAGATAGCCAGCGC and TATATAGCTAGCTCAAAGCGCATTTTTGGGCGTGTC. The resulting PCR product was digested with XhoI and NheI and cloned into pK91 (Belyy et al., 2016), a derivative of pESC-Ura in which expression is driven by the constitutive TEF1 promoter.

*Saccharomyces cerevisiae* MH272 haploid act1::LEU2 + ActinWT(His) (Belyy et al., 2015) was grown in synthetic defined yeast nitrogen base medium (Difco, #291940), containing glucose (2%) and supplemented, with uracil (2 µg/ml), tryptophan (2 µg/ml), and adenine (2 µg/ml). Yeast cells were transformed using the lithium acetate method (Gietz and Woods, 2002). Ceg14-dependent toxicity was assessed by a serial dilution spot assay as described previously (Belyy et al., 2015). Briefly, yeast cultures were normalized by OD600, subjected to fivefold serial dilutions, and spotted onto agar plates containing either glucose to repress Ceg14 expression or galactose with 0.05% glucose to induce expression from the galactose-responsive promoter.

Protein expression in yeast was analyzed by alkaline extraction of total cellular proteins as described previously (Kushnirov, 2000). Cells were incubated in 0.1 M NaOH for 5 min, collected, and resuspended directly in Laemmli sample buffer. Proteins were separated by SDS-PAGE, transferred to membranes, and analyzed by western blotting using an anti-Myc antibody (1:5,000; clone 9B11, #2276, Cell Signaling Technology).

### Cryogenic electron microscopy, data analysis and model building

Promptly after gel-filtration on the Superdex 200 increase 10/300GL column, 3 µl of the Ceg14-Actin-AnkJ complex (0.3 mg/ml) was applied twice onto a freshly glow-discharged copper R1.2/1.3 300 mesh grid (Quantifoil), with blotting for 3 s on both sides with blotting force 0 and plunge-frozen in liquid ethane-propane mixture using the Vitrobot Mark IV system (Thermo Fisher Scientific) at 13 °C and 100% humidity.

The cryo-EM dataset was collected using a Talos Arctica transmission electron microscope (Thermo Fisher Scientific) equipped with an XFEG at 200 kV using the automated data-collection software EPU. One image per hole with defocus range of -0.5 - -2.5 µm was collected with a Falcon 4i detector (Thermo Fisher Scientific). Image stacks with 38 frames were collected with a total exposure time of 5.2 sec and a total dose of 40 e-/Å^2^. 8,029 micrographs were used for particle processing in cryoSPARC (Punjani et al., 2017). Initial particle picking was performed with the general Topaz model (Bepler et al., 2019) on a subset of 3,034 micrographs, yielding 603,941 particles. Particles were extracted in 256-pixel boxes with 2-fold binning and subjected to two rounds of 2D classification. From the resulting classes, 80,138 particles were selected and used to train a dataset-specific Topaz model. Application of this model to the complete dataset yielded 939,307 particles, which were extracted in 256-pixel boxes with 2-fold binning and subjected to 2D classification. In parallel, blob picking was performed on the complete dataset, yielding 6,930,605 particles, which were extracted in 192-pixel boxes and subjected to 2D classification. Inspection of the resulting class averages revealed two major particle populations corresponding to the larger Ceg14-actin-AnkJ complex and the smaller Ceg14-AnkJ complex, which were processed independently.

For the **Ceg14-actin-AnkJ complex**, 126,173 particles selected from Topaz picking and 269,541 particles selected from blob picking were combined, re-extracted in 240-pixel boxes, and duplicate particles were removed. Following 2D classification, 211,455 particles were retained and subjected to ab initio reconstruction and non-uniform refinement, yielding a 3.4 Å reconstruction. After rebalancing the particle orientations, a further round of ab initio reconstruction and non-uniform refinement yielded a particle subset of 138,650 and a 3.5 Å reconstruction. Per-particle motion correction was subsequently performed, followed by non-uniform refinement with optimization of CTF parameters and per-particle defocus, improving the reconstruction to 3.2 Å. The particles were then exported to RELION (Scheres, 2012) and subjected to refinement with Blush regularization (Kimanius et al., 2024) and local-resolution filtering, resulting in the final 3.6 Å reconstruction.

For the **Ceg14-AnkJ complex**, 177,374 smaller particles selected from the Topaz-picked dataset and 448,096 smaller particles selected from the blob-picked dataset were combined, re-extracted in 192-pixel boxes, and duplicate particles were removed. After 2D classification, 363,011 particles were retained and subjected to ab initio reconstruction into three classes containing 143,302, 151,952 and 67,757 particles, respectively. The two classes corresponding to the Ceg14-AnkJ complex were combined, yielding 295,254 particles, and subjected to ab initio reconstruction followed by non-uniform refinement with optimization of CTF parameters, resulting in a 3.5 Å reconstruction. Following per-particle motion correction and an additional non-uniform refinement with CTF optimization, the reconstruction reached 3.4 Å. Particles were subsequently exported to RELION (Scheres, 2012) and refined with Blush regularization (Kimanius et al., 2024) followed by local-resolution filtering, yielding the final 3.6 Å reconstruction.

An initial model of AnkJ was generated using AlphaFold 3 (Abramson et al., 2024), whereas the structure of G-actin was taken from PDB 3HBT (Wang et al., 2010). Neither AlphaFold 3, nor Boltz-2 (Passaro et al., 2025) predictions of full-length Ceg14 did not align with the experimental density and was therefore not used directly. Instead, Ceg14 was predicted as separate fragments in Boltz-2, which produced models that could be reliably fitted into the corresponding regions of the cryo-EM map. The predicted AnkJ and Ceg14 fragments, together with the G-actin structure, were initially placed into the density by rigid-body fitting in UCSF Chimera (Pettersen et al., 2004). The resulting models were then interactively rebuilt and refined against the cryo-EM maps using ISOLDE (Croll, 2018), followed by final real-space refinement and model validation in Phenix (Liebschner et al., 2019).

Figures were prepared in UCSF Chimera.

### Sequence analysis

Ceg14 homologues were identified by BLASTP (Altschul et al., 1997) against the NCBI nr database (Sayers et al., 2022), using full-length *Legionella pneumophila* Philadelphia-1 Ceg14 as the query (UniProt Q5ZYD5, 666 aa; The UniProt Consortium, 2023), with E ≤ 10^-5^ and a maximum of 200 hits. Fragments and unrelated proteins were excluded by retaining only *Legionella* hits of ≥ 550 amino acids. This yielded 165 sequences from ten taxa, of which 137 were from *L. pneumophila*. These hits were aligned with MUSCLE v5.3 (Edgar, 2022), anchored on the reference so that only the 666 columns carrying a reference residue were kept, and displayed as a sequence logo generated with Logomaker (Tareen and Kinney, 2020). Per-position conservation was computed from an independent MAFFT v7.525 (--auto) alignment of the same sequences (Katoh and Standley, 2013) as the gap-free Shannon information content, as described (Shannon, 1948; Schneider and Stephens, 1990), scaled to [0,1] so that an invariant position scores 1. Five actin orthologues were retrieved from UniProt by their accession number (The UniProt Consortium, 2023) — human β-(P60709), γ-(P63261) and α-skeletal actin (P68133), *Dictyostelium discoideum* actin-1 (P07830) and *Saccharomyces cerevisiae* actin (P60010) — and aligned with MUSCLE v5.3 (Edgar, 2022).

### ATPase activity assay

ATP hydrolysis activity was quantified by measuring the release of inorganic phosphate (Pi) using the Malachite Green Phosphate Assay Kit (Sigma-Aldrich, Cat. No. MAK307). Where indicated, inorganic pyrophosphatase (PPiase) from baker’s yeast (Roche Diagnostics, Cat. No. 10108987001) was added after the reaction at a final concentration of 17 µg/mL and incubated for 5 min to convert pyrophosphate (PPi) to Pi. Thus, measurements performed without PPiase detected directly released Pi, whereas measurements performed with PPiase additionally detected PPi generated during ATP-to-AMP conversion. Reactions were performed in a total volume of 20 µL in TBS (20 mM Tris-HCl, pH 8.0, 150 mM NaCl) containing 1 mM ATP, 2 mM MgCl_2_ and 0.2 mg/mL Ceg14 or the indicated Ceg14 mutant. Where indicated, 2.2 µg of AnkJ, 3 µg of G-actin, Latrunculin A-stabilized G-actin, F-actin or phalloidin-stabilized F-actin, was added. D-(-)-3-phosphoglyceric acid disodium salt (3PG; Sigma-Aldrich, Cat. No. P8877) was added at 1 mM where indicated. Reactions were incubated for 25 min at 37 °C and subsequently diluted fourfold to reduce the ATP concentration to 0.25 mM, as suggested by the assay manual. Samples were mixed with freshly prepared Working Reagent (Reagent A : Reagent B, 100:1, v/v) in clear flat-bottom 96-well plates, and absorbance was measured at 620 nm using a SpectraMax ABS microplate reader (Molecular Devices).

### Molecular dynamics simulations

All systems were built from the cryo-EM structure of the Ceg14-actin-AnkJ complex: the ternary complex used the cryo-EM coordinates directly, the Ceg14-AnkJ complex was obtained by deleting the G-actin chain, and Ceg14 alone by deleting both AnkJ and actin. Systems were prepared in CHARMM-GUI Solution Builder (Jo et al., 2008; Lee et al., 2016) with the CHARMM36m force field (Huang et al., 2017) and TIP3P water (Jorgensen et al., 1983), placed in a cubic box and neutralized with Na⁺ and Cl⁻ ions to 0.15 M NaCl (Ceg14 alone: 14.0 nm box, 258783 atoms, 241 Na⁺, 236 Cl⁻, 83024 waters; Ceg14-AnkJ: 13 nm box, 206,293 atoms, 189 Na⁺, 182 Cl⁻, 64193 waters; Ceg14-actin-AnkJ: 15.5 nm box, 352380 atoms, 334 Na⁺, 314 Cl⁻, 110854 waters). All simulations were performed in GROMACS (Abraham et al., 2015): version 2024.4 for minimization and equilibration, and version 2025.1 for production on NVIDIA H100. Each system was first energy-minimized by steepest descent (to Fmax < 1000 kJ mol⁻¹ nm⁻¹) with the protein held in place by position restraints on backbone (400 kJ mol⁻¹ nm⁻²) and side-chain (40 kJ mol⁻¹ nm⁻²) heavy atoms, then equilibrated for 125 ps under the same restraints at 310 K in the NVT ensemble (1 fs time step, v-rescale thermostat (Bussi et al., 2007), τᵀ = 1.0 ps, protein and solvent coupled separately, starting velocities drawn from a Maxwell–Boltzmann distribution at 310 K). The restraints were then removed and production runs carried out in the NPT ensemble at 310 K and 1 bar with a 2 fs time step, using the same thermostat together with the C-rescale barostat (Bernetti and Bussi, 2020) (isotropic, τᴾ = 5.0 ps, compressibility 4.5 × 10⁻⁵ bar⁻¹). Bonds to hydrogen were constrained with LINCS (Hess et al., 1997), long-range electrostatics were treated by particle-mesh Ewald (Essmann et al., 1995) with a 1.2 nm real-space cutoff, van der Waals interactions were switched off between 1.0 and 1.2 nm using the Verlet neighbor scheme (updated every 20 steps), and center-of-mass motion was removed every 100 steps. Each system was run in three replicates, all started from the same equilibrated structure but with different random starting velocities (gen_seed = −1 at the production grompp step); each replicate was simulated for 400 ns. Analyses were performed with the GROMACS 2025.1 analysis tools (Abraham et al., 2015) and MDAnalysis 2.10 (Michaud-Agrawal et al., 2011; Gowers et al., 2016), using NumPy (Harris et al., 2020) and Matplotlib (Hunter, 2007) for averaging and plotting. Every quantity was calculated separately for each of the three replicates of a system and is reported as the mean of the three, with the spread between them given as the standard deviation. Per-residue root-mean-square fluctuations (RMSF) of the Ceg14 Cα atoms were obtained with *gmx rmsf* over the final 100 ns of each trajectory (300–400 ns), after fitting the trajectory on the Cα atoms of all protein chains to remove overall translation and rotation. The distance between the mass-weighted centroid of the three catalytic Cα atoms Ser527, His571 and Glu575 and the mass-weighted centroid of the Arg441, Arg443 and Lys456 Cα atoms was measured with MDAnalysis.

## Author Contributions

Conceptualization (D.S., A.B.), Data Curation (D.S., A.B.), Formal Analysis (D.S., A.B.), Funding Acquisition (A.B.), Investigation (D.S., M.S.E., A.B.), Methodology (D.S., S.J.M., G.N.S., A.B.), Project Administration (A.B.), Resources (S.J.M., G.N.S., A.B.), Software (D.S., S.J.M.), Supervision (S.J.M., G.N.S., A.B.), Validation (D.S., A.B.), Visualization (D.S., A.B.), Writing – Original Draft Preparation (D.S., M.S.E, A.B.), Writing – Review & Editing (all authors).

## Acknowledgements

We thank Michiel Punter for maintaining the cryo-EM computing cluster, Artem Stetsenko for help with cryo-EM data collection, Laura Darkhan for help in cloning of Ceg14 and AnkJ in yeast vectors, and members of the Marrink, Schroeder and Belyy labs for helpful discussions. Cryo-EM data were collected at the electron microscopy facility of the University of Groningen. This work was supported by ERC Starting Grant “ACTIN in ACTION” #101219748 and Dutch Research Council NWO OCENW.M.24.262. MD simulations and structural predictions were performed on NWO Snellius cluster, supported by the computational grant EINF-19040.

## Data availability

Cryo-EM reconstructions for the Ceg14-actin-AnkJ and Ceg14-AnkJ complexes have been deposited in the Electron Microscopy Data Bank under accession numbers EMD-59299 and EMD-59298, respectively. The corresponding molecular models have been deposited at the wwPDB with accession codes 33AH and 33AG. The raw data generated during the current study are available from the corresponding author on request.

The authors have declared no competing interest.

## Supporting information

**Supplementary Figure S1.**
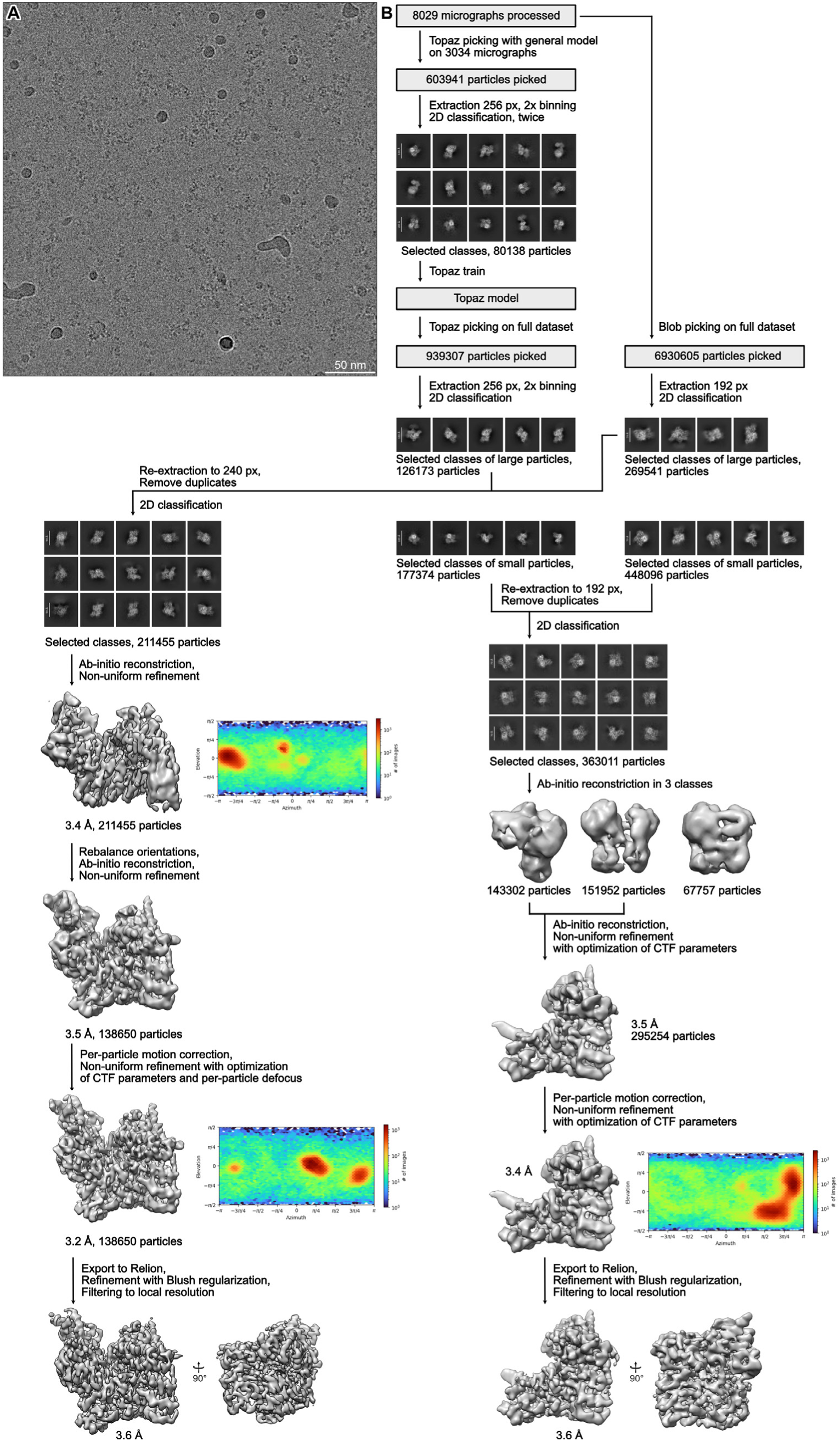
Data processing of the cryo-EM dataset. (A) Exemplary cryo-EM micrograph. (B) Processing pipeline, showing examples of 2D classes and intermediate 3D reconstructions with their angular distributions.

**Supplementary Figure S2.**
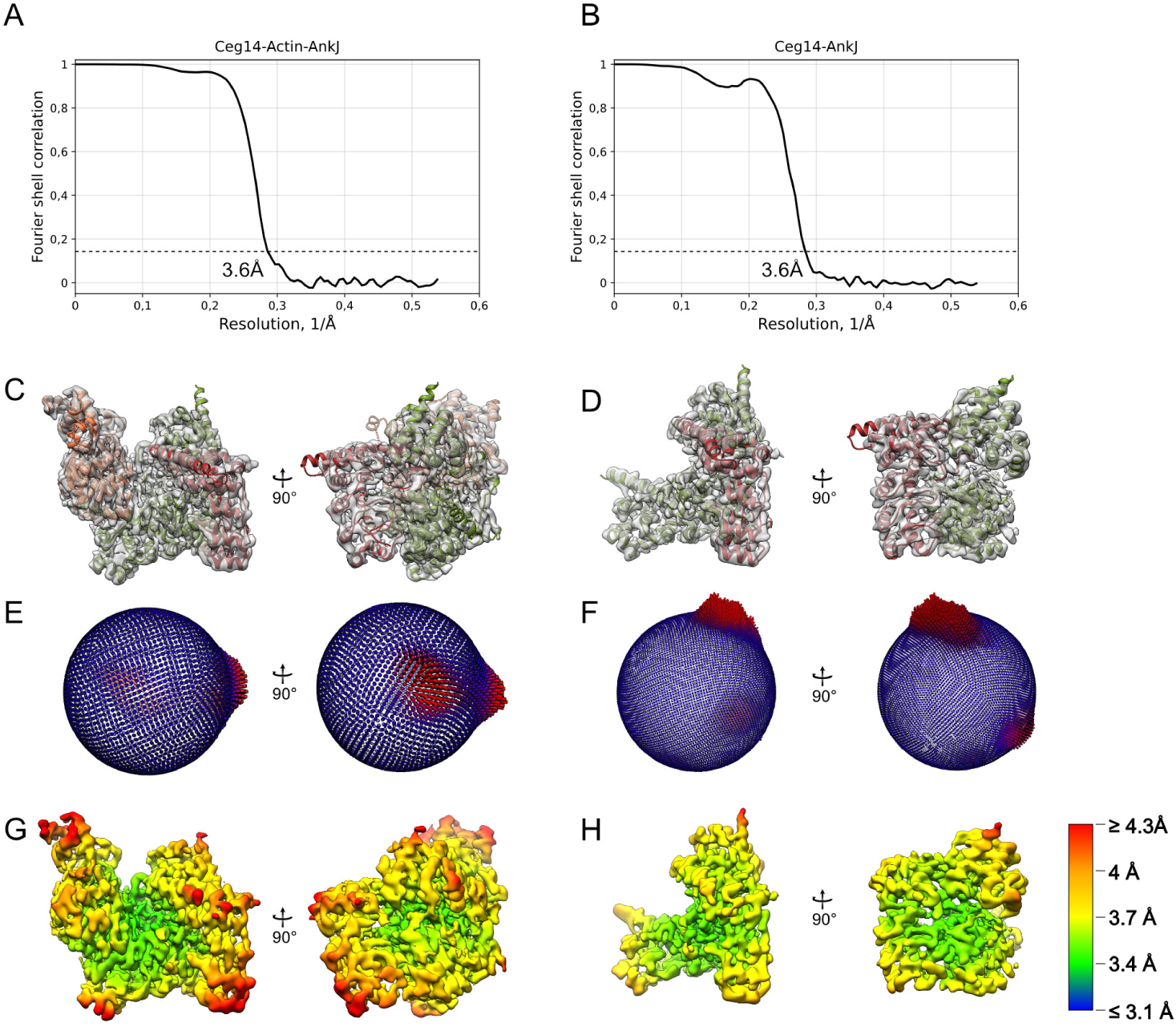
Quality assessment of cryo-EM reconstructions. Fourier shell correlation curves of the final masked volumes of the Ceg14-actin-AnkJ (A) and Ceg14-AnkJ (B) reconstructions. Fitting of the Ceg14-actin-AnkJ (C) and Ceg14-AnkJ (D) structures into the corresponding cryo-EM densities. Angular distribution of the Ceg14-actin-AnkJ (E) and Ceg14-AnkJ (F) reconstructions. Local resolution of the Ceg14-actin-AnkJ (G) and Ceg14-AnkJ (H) reconstructions.

**Supplementary Figure S3.**
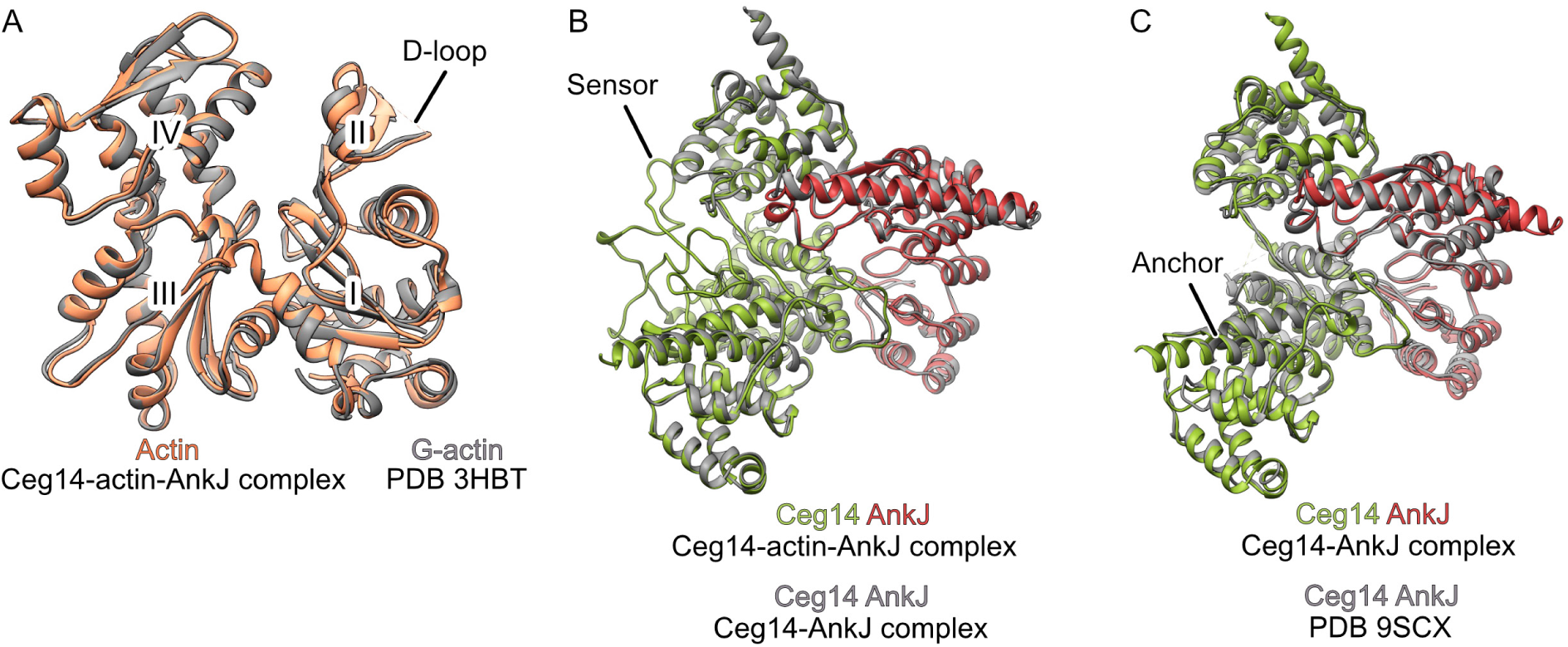
Comparisons of cryo-EM structures. (A) Alignment of the crystal structure of G-actin (PDB 3HBT) to actin from the Ceg14-actin-AnkJ complex. Actin subdomains I to IV are labelled with roman numerals. (B) Alignment of structures of Ceg14 and AnkJ from the Ceg14-actin-AnkJ and the Ceg14-AnkJ complexes. (C) Structural alignment of Ceg14 and AnkJ from the Ceg14-AnkJ complex to the previously published crystal structure (PDB 9SCX).

**Supplementary Figure S4.**
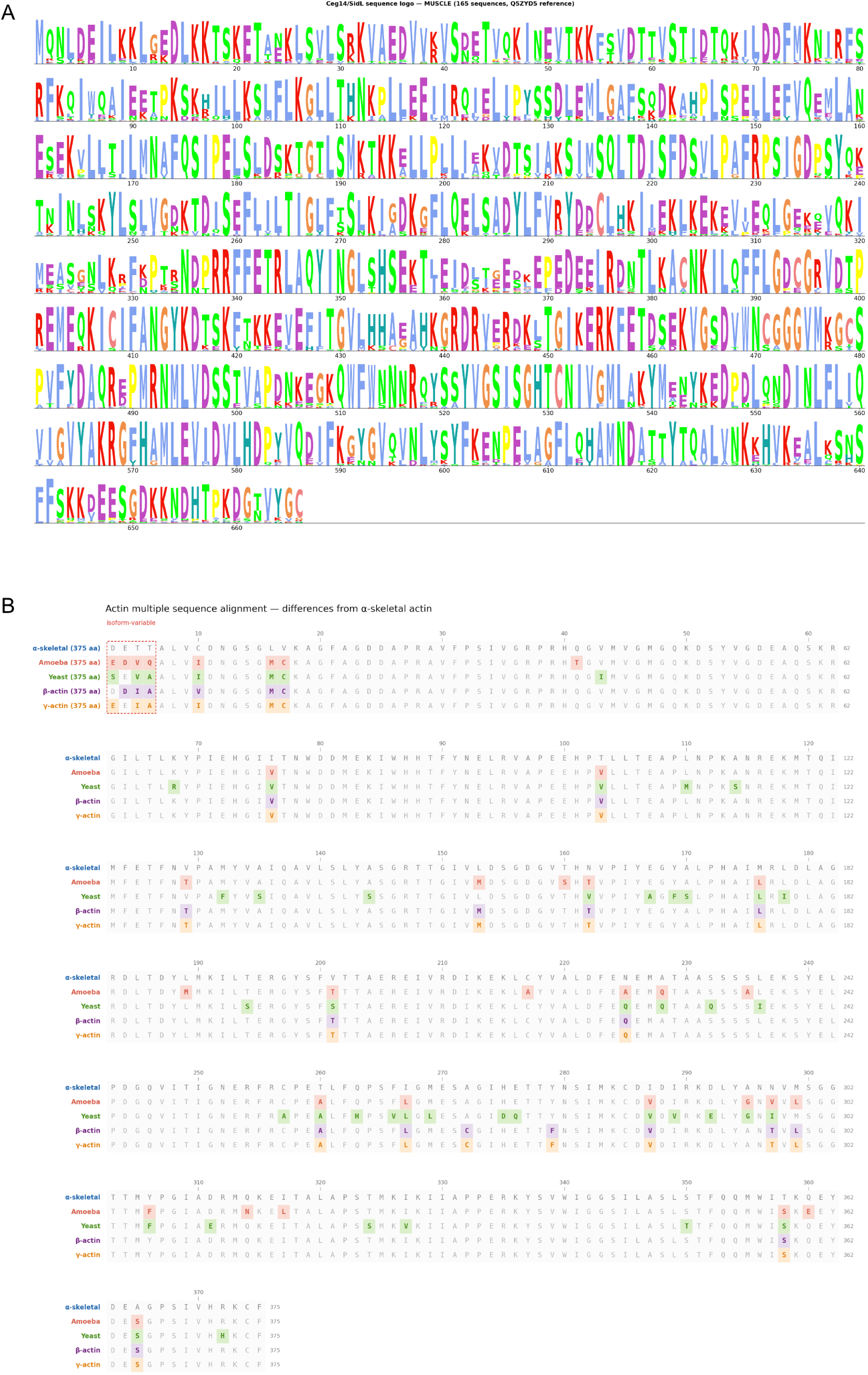
Sequence conservation of Ceg14 and Actin. (A) Conservation of amino acids in Ceg14 among 165 unique sequences. (B) Sequence comparison of yeast, amoeba and human actins.

**Supplementary Table S1:** Cryo-EM data collection, refinement, and validation statistics.

|  |  |  |
| --- | --- | --- |
|  | Ceg14-AnkJ complex | Ceg14-actin-AnkJ complex |
| Microscope | Talos Arctica |  |
| Voltage (kV) | 200 |  |
| Defocus range (μm) | -0.5 to -2.5 |  |
| Camera | Falcon 4i |  |
| Pixel size (Å) | 0.93 |  |
| Total electron dose (e/Å<sup>2</sup>) | 40 |  |
| Exposure time (s) | 5.2 |  |
| Frames per movie | 38 |  |
| Number of movies | 8029 (11273 collected) |  |
| 3D Refinement |  |  |
| Number of particles | 295254 | 138650 |
| Final map pixel size | 0.93 | 0.93 |
| Final resolution (Å) | 3.6 | 3.6 |
| Atomic model statistics |  |  |
| Non-hydrogen atoms | 5680 | 9436 |
| Number of protein chains | 2 | 3 |
| Molprobity score | 1.16 | 1.16 |
| Rama distribution Z-score | -2.31 ± 0.24 | -1.08 ± 0.23 |
| Clashscore | 2.01 | 2.33 |
| Bond RMSD (Å) | 0.002 | 0.003 |
| Angle RMSD (°) | 0.551 | 0.632 |
| Cβ deviations >0.25Å | 0 | 0 |
| Poor rotamers (%) | 0 | 0 |
| Favored rotamers (%) | 99.84 | 99.81 |
| Ramachandran favored (%) | 96.86 | 97.18 |
| Ramachandran allowed (%) | 3.14 | 2.82 |
| Ramachandran outliers (%) | 0 | 0 |
| Missing fragments | Ceg14: 1-70, 414-528, 638-666.<br>AnkJ: 1-5, 257-269 | Actin: 1-4, 40-50.<br>Ceg14: 1-71, 641-666<br>AnkJ: 1-5, 257-269 |

**Supplementary Table S2:** Data points Figure 1 A.

| Conditions | Repeat 1 | Repeat 2 | Repeat 3 |
| --- | --- | --- | --- |
| Ceg14 WT + PPiase | 0,294806 | -1,60038 | 0,08423 |
| Ceg14 WT + G-actin + PPiase | 422,9447 | 288,3031 | 303,925 |
| Ceg14 WT + G-actin-LatA + PPiase | 414,5479 | 317,106 | 303,7297 |
| Ceg14 WT + F-actin-PHD + PPIase | 23,2897 | 27,03795 | 11,75014 |
| Ceg14 WT + G-actin-LatA | 4,464211 | 4,169404 | 13,56109 |
| Ceg14 E575A + G-actin-LatA + PPIase | -0,75807 | 3,621907 | -1,05288 |

**Figure 1 B**
| Conditions | Repeat 1 | Repeat 2 | Repeat 3 |
| --- | --- | --- | --- |
| Ceg14 WT alone + PPIase | -2,23210525 | 1,431916578 | 1,853068513 |
| Ceg14 WT + AnkJ + PPIase | 5,011708024 | 9,433803339 | -0,46326713 |
| Ceg14 WT + G-actin-LatA + AnkJ + PPIase | 21,9420158 | 27,16429979 | 33,65003959 |
| Ceg14 WT + G-actin-LatA + AnkJ | 24,46892741 | 18,44645474 | 29,18582908 |
| Ceg14 E575A + G-actin-LatA + AnkJ + PPIase | -1,09499503 | 1,768838126 | 9,854955274 |
| Ceg14 E575A + G-actin-LatA + AnkJ | 5,011708024 | 1,052879837 | 3,8745978 |

**Figure 1 D**
| Conditions | Repeat 1 | Repeat 2 | Repeat 3 |
| --- | --- | --- | --- |
| Ceg14 WT + PPIase | -0,48432473 | 1,9372989 | 1,300306599 |
| Ceg14 WT + 3PG + PPIase | 1,295042199 | 1,784631324 | 2,563762403 |
| Ceg14 WT + G-actin-LatA + PPIase | 388,3811756 | 340,3436829 | 322,7689904 |
| Ceg14 WT + G-actin-LatA + 3PG + PPIase | 541,9013714 | 645,5469935 | 430,4447749 |
| Ceg14 WT + G-actin-LatA + 3PG + AnkJ + PPIase | 22,63691649 | 31,77591348 | 26,32199592 |
| Ceg14 WT + G-actin-LatA + 3PG + AnkJ | 18,27799397 | 30,59668806 | 26,42728391 |

**Figure 2 D**
| Conditions | Repeat 1 | Repeat 2 | Repeat 3 |
| --- | --- | --- | --- |
| Ceg14 WT + G-actin-LatA + PPIase | 422,9447374 | 447,0611209 | 343,3704355 |
| Ceg14 V632A L636A + G-actin-LatA + PPIase | -1,55826216 | 3,579791446 | -6,65420057 |
| Ceg14 R446E + G-actin-LatA + PPIase | 29,0594835 | 21,22605751 | 8,549384276 |
| Ceg14 D485K D618K + G-actin-LatA + PPIase | -0,71595829 | -0,54749752 | -3,49556106 |
| Ceg14 R441S R443S K456S + G-actin-LatA + PPIase | 27,50122134 | 20,97336635 | 19,16241303 |
| Ceg14 S221G F222G + G-actin-LatA + PPIase | 20,67856 | 24,42681222 | 17,35145971 |

## References

Abraham MJ et al. 2015. GROMACS: High performance molecular simulations through multi-level parallelism from laptops to supercomputers. SoftwareX 1-2:19–25. doi:10.1016/j.softx.2015.06.001.

Abramson J et al. 2024. Accurate structure prediction of biomolecular interactions with AlphaFold 3. Nature 630:493–500. doi:10.1038/s41586-024-07487-w.

Aktories K, Lang AE, Schwan C, Mannherz HG. 2011. Actin as target for modification by bacterial protein toxins. FEBS J 278:4526–4543. doi:10.1111/j.1742-4658.2011.08113.x.

Altschul SF et al. 1997. Gapped BLAST and PSI-BLAST: a new generation of protein database search programs. Nucleic Acids Res 25:3389–3402. doi:10.1093/nar/25.17.3389.

Belmont LD, Drubin DG. 2001. Actin structure-function relationships revealed by yeast molecular genetics. Results Probl Cell Differ 32:103–121. doi:10.1007/978-3-540-46560-7_8.

Belyy A et al. 2015. Roles of Asp179 and Glu270 in ADP-ribosylation of actin by Clostridium perfringens iota toxin. PLoS One 10:e0145708. doi:10.1371/journal.pone.0145708.

Belyy A et al. 2016. Ribosomal protein Rps26 influences 80S ribosome assembly in Saccharomyces cerevisiae. mSphere 1:e00109–15. doi:10.1128/mSphere.00109-15.

Belyy A et al. 2018. The extreme C terminus of the Pseudomonas aeruginosa effector ExoY is crucial for binding to its eukaryotic activator, F-actin. J Biol Chem 293:19785–19796. doi:10.1074/jbc.RA118.003784.

Belyy A et al. 2021. Mechanism of actin-dependent activation of nucleotidyl cyclase toxins from bacterial human pathogens. Nat Commun 12:6628. doi:10.1038/s41467-021-26889-2.

Belyy A et al. 2022. Mechanism of threonine ADP-ribosylation of F-actin by a Tc toxin. Nat Commun 13:4202. doi:10.1038/s41467-022-31836-w.

Bepler T et al. 2019. Positive-unlabeled convolutional neural networks for particle picking in cryo-electron micrographs. Nat Methods 16:1153–1160. doi:10.1038/s41592-019-0575-8.

Bernetti M, Bussi G. 2020. Pressure control using stochastic cell rescaling. J Chem Phys 153:114107. doi:10.1063/5.0020514.

Black JJ et al. 2026. The Legionella pneumophila effector SidL is an adenylyltransferase that modifies the glycolytic intermediate 3-phosphoglycerate. Mol Cell. doi:10.1016/j.molcel.2026.07.007.

Boamah DK, Zhou G, Ensminger AW, O’Connor TJ. 2017. From many hosts, one accidental pathogen: the diverse protozoan hosts of Legionella. Front Cell Infect Microbiol 7:477. doi:10.3389/fcimb.2017.00477.

Bussi G, Donadio D, Parrinello M. 2007. Canonical sampling through velocity rescaling. J Chem Phys 126:014101. doi:10.1063/1.2408420.

Campodonico EM, Chesnel L, Roy CR. 2005. A yeast genetic system for the identification and characterization of substrate proteins transferred into host cells by the Legionella pneumophila Dot/Icm system. Mol Microbiol 56:918–933. doi:10.1111/j.1365-2958.2005.04595.x.

Carlier MF. 1990. Actin polymerization and ATP hydrolysis. Adv Biophys 26:51–73. doi:10.1016/0065-227X(90)90007-G.

Chen TT et al. 2026. Structure and mechanism of an actin-dependent bacterial phosphoryl AMPylase. Nat Chem Biol 22:152–162. doi:10.1038/s41589-025-01945-w.

Croll TI. 2018. ISOLDE: a physically realistic environment for model building into low-resolution electron-density maps. Acta Crystallogr D Struct Biol 74:519–530. doi:10.1107/S2059798318002425.

Edgar RC. 2022. Muscle5: high-accuracy alignment ensembles enable unbiased assessments of sequence homology and phylogeny. Nat Commun 13:6968. doi:10.1038/s41467-022-34630-w.

Essmann U et al. 1995. A smooth particle mesh Ewald method. J Chem Phys 103:8577–8593. doi:10.1063/1.470117.

Estes JE, Selden LA, Gershman LC. 1981. Mechanism of action of phalloidin on the polymerization of muscle actin. Biochemistry 20:708–712. doi:10.1021/bi00507a006.

Fontana MF et al. 2011. Secreted bacterial effectors that inhibit host protein synthesis are critical for induction of the innate immune response to virulent Legionella pneumophila. PLoS Pathog 7:e1001289. doi:10.1371/journal.ppat.1001289.

Franco IS, Shohdy N, Shuman HA. 2012. The Legionella pneumophila effector VipA is an actin nucleator that alters host cell organelle trafficking. PLoS Pathog 8:e1002546. doi:10.1371/journal.ppat.1002546.

Fraser DW et al. 1977. Legionnaires’ disease: description of an epidemic of pneumonia. N Engl J Med 297:1189–1197. doi:10.1056/NEJM197712012972201.

Fu J et al. 2024. Legionella maintains host cell ubiquitin homeostasis by effectors with unique catalytic mechanisms. Nat Commun 15:5953. doi:10.1038/s41467-024-50311-2.

Funk J et al. 2019. Profilin and formin constitute a pacemaker system for robust actin filament growth. eLife 8:e50963. doi:10.7554/eLife.50963.

Gietz RD, Woods RA. 2002. Transformation of yeast by lithium acetate/single-stranded carrier DNA/polyethylene glycol method. Methods Enzymol 350:87–96. doi:10.1016/S0076-6879(02)50957-5.

Gowers RJ et al. 2016. MDAnalysis: a Python package for the rapid analysis of molecular dynamics simulations. Proc 15th Python Sci Conf:98–105. doi:10.25080/Majora-629e541a-00e.

Guan H et al. 2026. Molecular basis of host ATP level modulation by actin-dependent secreted bacterial ATPase and its metaeffector. Nat Commun 17:7846. doi:10.1038/s41467-026-74513-y.

Gülke I et al. 2001. Characterization of the enzymatic component of the ADP-ribosyltransferase toxin CDTa from Clostridium difficile. Infect Immun 69:6004–6011. doi:10.1128/IAI.69.10.6004-6011.2001.

Guo Z, Stephenson R, Qiu J, Zheng S, Luo ZQ. 2014. A Legionella effector modulates host cytoskeletal structure by inhibiting actin polymerization. Microbes Infect 16:225–236. doi:10.1016/j.micinf.2013.11.007.

Harris CR et al. 2020. Array programming with NumPy. Nature 585:357–362. doi:10.1038/s41586-020-2649-2.

He C et al. 2025. Modulation of host ATP levels by secreted bacterial effectors. Nat Commun 16:4675. doi:10.1038/s41467-025-60046-3.

Heidtman M, Chen EJ, Moy MY, Isberg RR. 2009. Large-scale identification of Legionella pneumophila Dot/Icm substrates that modulate host cell vesicle trafficking pathways. Cell Microbiol 11:230–248. doi:10.1111/j.1462-5822.2008.01249.x.

Hess B et al. 1997. LINCS: a linear constraint solver for molecular simulations. J Comput Chem 18:1463–1472. doi:10.1002/(SICI)1096-987X(199709)18:12<1463::AID-JCC4>3.0.CO;2-H.

Hiyoshi H et al. 2011. VopV, an F-actin-binding type III secretion effector, is required for Vibrio parahaemolyticus-induced enterotoxicity. Cell Host Microbe 10:401–409. doi:10.1016/j.chom.2011.08.014.

Huang J et al. 2017. CHARMM36m: an improved force field for folded and intrinsically disordered proteins. Nat Methods 14:71–73. doi:10.1038/nmeth.4067.

Hunter JD. 2007. Matplotlib: a 2D graphics environment. Comput Sci Eng 9:90–95. doi:10.1109/MCSE.2007.55.

Isberg RR, O’Connor TJ, Heidtman M. 2009. The Legionella pneumophila replication vacuole: making a cosy niche inside host cells. Nat Rev Microbiol 7:13–24. doi:10.1038/nrmicro1967.

Jo S, Kim T, Iyer VG, Im W. 2008. CHARMM-GUI: a web-based graphical user interface for CHARMM. J Comput Chem 29:1859–1865. doi:10.1002/jcc.20945.

Jorgensen WL, Chandrasekhar J, Madura JD, Impey RW, Klein ML. 1983. Comparison of simple potential functions for simulating liquid water. J Chem Phys 79:926–935. doi:10.1063/1.445869.

Joseph AM et al. 2020. The Legionella pneumophila metaeffector Lpg2505 (MesI) regulates SidI-mediated translation inhibition and novel glycosyl hydrolase activity. Infect Immun 88:e00853–19. doi:10.1128/IAI.00853-19.

Joseph AM, Shames SR. 2021. Affecting the Effectors: Regulation of Legionella pneumophila Effector Function by Metaeffectors. Pathogens 10(2):108. doi: 10.3390/pathogens10020108

Katoh K, Standley DM. 2013. MAFFT multiple sequence alignment software version 7: improvements in performance and usability. Mol Biol Evol 30:772–780. doi:10.1093/molbev/mst010.

Kimanius D et al. 2024. Data-driven regularization lowers the size barrier of cryo-EM structure determination. Nat Methods 21:1216–1221. doi:10.1038/s41592-024-02304-8.

Kushnirov VV. 2000. Rapid and reliable protein extraction from yeast. Yeast 16:857–860. doi:10.1002/1097-0061(20000630)16:9<857::AID-YEA561>3.0.CO;2-B.

Lee J et al. 2016. CHARMM-GUI Input Generator for NAMD, GROMACS, AMBER, OpenMM, and CHARMM/OpenMM simulations using the CHARMM36 additive force field. J Chem Theory Comput 12:405–413. doi:10.1021/acs.jctc.5b00935.

Liebschner D et al. 2019. Macromolecular structure determination using X-rays, neutrons and electrons: recent developments in Phenix. Acta Crystallogr D Struct Biol 75:861–877. doi:10.1107/S2059798319011471.

Liu X, Shin S. 2019. Viewing Legionella pneumophila pathogenesis through an immunological lens. J Mol Biol 431:4321–4344. doi:10.1016/j.jmb.2019.07.028.

Liu Y et al. 2017. A Legionella effector disrupts host cytoskeletal structure by cleaving actin. PLoS Pathog 13:e1006186. doi:10.1371/journal.ppat.1006186.

Liu Y, Liu Y, Luo ZQ. 2025. Legionella pneumophila modulates the host cytoskeleton by an effector of transglutaminase activity. mLife 4:232–248. doi:10.1002/mlf2.70013.

Lockwood DC, Amin H, Costa TRD, Schroeder GN. 2022. The Legionella pneumophila Dot/Icm type IV secretion system and its effectors. Microbiology 168:001187. doi:10.1099/mic.0.001187.

Machtens DA et al. 2026. Crystal structure of the Legionella pneumophila effector SidL (Lpg0437) in complex with its metaeffector LegA11 (Lpg0436). Virulence 17:2646775. doi:10.1080/21505594.2026.2646775.

McDade JE et al. 1977. Legionnaires’ disease: isolation of a bacterium and demonstration of its role in other respiratory disease. N Engl J Med 297:1197–1203. doi:10.1056/NEJM197712012972202.

Michaud-Agrawal N et al. 2011. MDAnalysis: a toolkit for the analysis of molecular dynamics simulations. J Comput Chem 32:2319–2327. doi:10.1002/jcc.21787.

Niedzialkowska E et al. 2024. Stabilization of F-actin by Salmonella effector SipA resembles the structural effects of inorganic phosphate and phalloidin. Structure 32:725–738.e8. doi:10.1016/j.str.2024.02.022.

Passaro S et al. 2025. Boltz-2: Towards Accurate and Efficient Binding Affinity Prediction. bioRxiv. doi:10.1101/2025.06.14.659707.

Peisker K et al. 2008. Ribosome-associated complex binds to ribosomes in close proximity of Rpl31 at the exit of the polypeptide tunnel in yeast. Mol Biol Cell 19:5279–5288. doi:10.1091/mbc.E08-06-0661.

Pettersen EF et al. 2004. UCSF Chimera - a visualization system for exploratory research and analysis. J Comput Chem 25:1605–1612. doi:10.1002/jcc.20084.

Pollard DT. 2016. Actin and Actin-Binding Proteins. Cold Spring Harb Perspect Biol. 2016 Aug 1;8(8):a018226. doi: 10.1101/cshperspect.a018226.

Pospich S, Merino F, Raunser S. 2020. Structural effects and functional implications of phalloidin and jasplakinolide binding to actin filaments. Structure 28:437–449.e5. doi:10.1016/j.str.2020.01.014.

Price C, Jones S, Mihelcic M, Santic M, Abu Kwaik Y. 2020. Paradoxical pro-inflammatory responses by human macrophages to an amoebae host-adapted Legionella effector. Cell Host Microbe 27:571–584.e7. doi:10.1016/j.chom.2020.03.003.

Punjani A, Rubinstein JL, Fleet DJ, Brubaker MA. 2017. cryoSPARC: algorithms for rapid unsupervised cryo-EM structure determination. Nat Methods 14:290–296. doi:10.1038/nmeth.4169.

Sayers EW et al. 2022. Database resources of the National Center for Biotechnology Information. Nucleic Acids Res 50:D20–D26. doi:10.1093/nar/gkab1112.

Scheres SHW. 2012. RELION: implementation of a Bayesian approach to cryo-EM structure determination. J Struct Biol 180:519–530. doi:10.1016/j.jsb.2012.09.006.

Schneider TD, Stephens RM. 1990. Sequence logos: a new way to display consensus sequences. Nucleic Acids Res 18:6097–6100. doi:10.1093/nar/18.20.6097.

Shannon CE. 1948. A mathematical theory of communication. Bell Syst Tech J 27:379–423, 623-656. doi:10.1002/j.1538-7305.1948.tb01338.x

Tan Y, Luo ZQ. 2011. Legionella pneumophila SidD is a deAMPylase that modifies Rab1. Nature 475:506–509. doi:10.1038/nature10307.

Tareen A, Kinney JB. 2020. Logomaker: beautiful sequence logos in Python. Bioinformatics 36:2272–2274. doi:10.1093/bioinformatics/btz921.

The UniProt Consortium. 2023. UniProt: the Universal Protein Knowledgebase in 2023. Nucleic Acids Res 51:D523–D531. doi:10.1093/nar/gkac1052.

Tsuge H et al. 2008. Structural basis of actin recognition and arginine ADP-ribosylation by Clostridium perfringens iota-toxin. Proc Natl Acad Sci USA 105:7399–7404. doi:10.1073/pnas.0801215105.

Urbanus ML et al. 2016. Diverse mechanisms of metaeffector activity in an intracellular bacterial pathogen, Legionella pneumophila. Mol Syst Biol 12:893. doi:10.15252/msb.20167381.

Wang H, Robinson RC, Burtnick LD. 2010. The structure of native G-actin. Cytoskeleton 67:456–465. doi:10.1002/cm.20458.

Wang T et al. 2024. Legionella effector LnaB is a phosphoryl-AMPylase that impairs phosphosignalling. Nature 631:393–401. doi:10.1038/s41586-024-07573-z.

Yarmola EG et al. 2000. Actin-latrunculin A structure and function: differential modulation of actin-binding protein function by latrunculin A. J Biol Chem 275:28120–28127. doi:10.1074/jbc.M004253200.

Zhang Q et al. 2023. Membrane-dependent actin polymerization mediated by the Legionella pneumophila effector protein MavH. PLoS Pathog 19:e1011512. doi:10.1371/journal.ppat.1011512.

Zhang Q et al. 2026. Legionella effector RavH is a tandem WH2-like domain-containing PI(3)P-dependent actin nucleator. Life Sci Alliance. doi:10.26508/lsa.202603675.

